# Characterization of endogenous Rubus yellow net virus in raspberries

**DOI:** 10.1101/2021.06.17.448838

**Authors:** Thien Ho, Janet C. Broome, Jason P. Buhler, Wendy O’Donovan, Tongyan Tian, Alfredo Diaz-Lara, Robert R. Martin, Ioannis E. Tzanetakis

## Abstract

Rubus yellow net virus (RYNV) belongs to genus *Badnavirus*. Badnavirids are found in plants as endogenous, inactive sequences, and/or in episomal (infectious and active) forms. To assess the state of RYNV infections, we sequenced the genomes of various *Rubus* cultivars and mined eight additional published whole genome sequencing datasets. Sequence analysis revealed the presence of a diverse array of endogenous RYNV (endoRYNV) sequences that differ significantly in their structure; some lineages have nearly complete, yet non-functional genomes whereas others have rudimentary, small sequence fragments. We developed assays to genotype the six main endoRYNV lineages as well as the only known episomal lineage in commercial *Rubus*. This study discloses the widespread presence of endoRYNVs in commercial raspberries, likely because breeding programs have been using a limited pool of germplasm that harbored endoRYNVs.

## 1. Introduction

Raspberry is an economically important crop with global production in 2018 being over 820,000 tons grown in 125,000 hectares in all continents except Antarctica (“Production of raspberries in 2018”. United Nations, Corporate Statistical Database (FAOSTAT) 2019, retrieved January 14, 2021). Commercial breeding for red raspberry (*Rubus idaeus*) began about 200 years ago, and most of the currently available cultivars share the same germplasm pedigree dating back to the late 1800s and early 1900s (Jennings, 2018).

More than 40 virus species are known to infect *Rubus*; yet Rubus yellow net virus (RYNV) is one of only two badnaviruses known to infect the genus (Diaz-Lara et al., 2015; Shahid et al., 2017). The genus *Badnavirus*, family *Caulimoviridae*, includes viruses that have an inactive, endogenous form and an infectious, episomal form. *Banana streak OL virus* (BSOLV), *Banana streak GF virus* (BSGFV), and *Banana streak IM virus* (BSIMV) integrants have been shown to reactivate from an inactive integrated counterpart (Chabannes et al., 2013; Gayral et al., 2008; Ndowora et al., 1999), whereas no other badnavirid is known to reactivate from integrated sequences (reviewed by Bhat et al., 2016).

RYNV is a component of raspberry mosaic, an important disease first described in the 1920s (Bennett, 1927; Stace-Smith, 1955). The virus infects all red raspberry and most blackberry and hybrid berry cultivars in North America and Europe (Stace-Smith and Jones, 1987a) and can reduce yield from 30-75% in the first year and up to 15% in subsequent years in mixed infections with black raspberry necrosis virus (Stace-Smith and Jones, 1987b). Partial RYNV sequences were first obtained at the turn of the century (GenBank accession number AF468454; Jones et al., 2002). The plant used in the Jones et al. (2002) study had virus-like symptoms and bacilliform particles were observed under the electron microscope. The first RYNV genome (RYNV-Ca, GenBank accession number KF241951), assembled from two PCR amplicons, was obtained by Kalischuk et al. (2008). Another genome (RYNV-BS, KM078034, Diaz-Lara et al., 2015) was sequenced from red raspberry ‘Baumforth’s Seedling A’ using DNA from rolling circle amplification. Since then, several RYNV sequences were published using small RNA (Rajamäki et al., 2019) or whole genome sequencing (MN245240).

Diaz-Lara (2016) observed that red raspberry plants, supposedly free of RYNV based on aphid or graft transmission onto *R. occidentalis* ‘Munger’ indicator, yielded positive results when indexed by PCR-based assays. Moreover, those plants were tested positive for RYNV by PCR even after heat therapy and meristem-tip culture for virus elimination. It was demonstrated that RYNV integrates into the red raspberry genome (Diaz-Lara et al., 2020), but no further analysis was conducted for the reported endogenous RYNV (endoRYNV) sequences. In this study, multiple cultivars were assayed to determine the prevalence of endoRYNV and the lineages identified were validated and characterized in-depth.

## 2. Materials and methods

### 2.1. Plant material

Twenty-five raspberry cultivars maintained as tissue culture plantlets in Watsonville, California were used in the study (Table 1). For ‘Baumforth’s Seedling A’, an additional mature plant was obtained from Corvallis, Oregon with the RYNV-BS (Diaz-Lara et al., 2015; Table 1) and used as a positive control for the episomal form, hereafter referred to as epiRYNV-BS. In addition, the genome of 75 proprietary red raspberry and 100 proprietary blackberry cultivars were sequenced and assayed for integration of RYNV-BS and the episomal form of the virus but their identity is not provided to protect intellectual property rights.

**Table 1.**
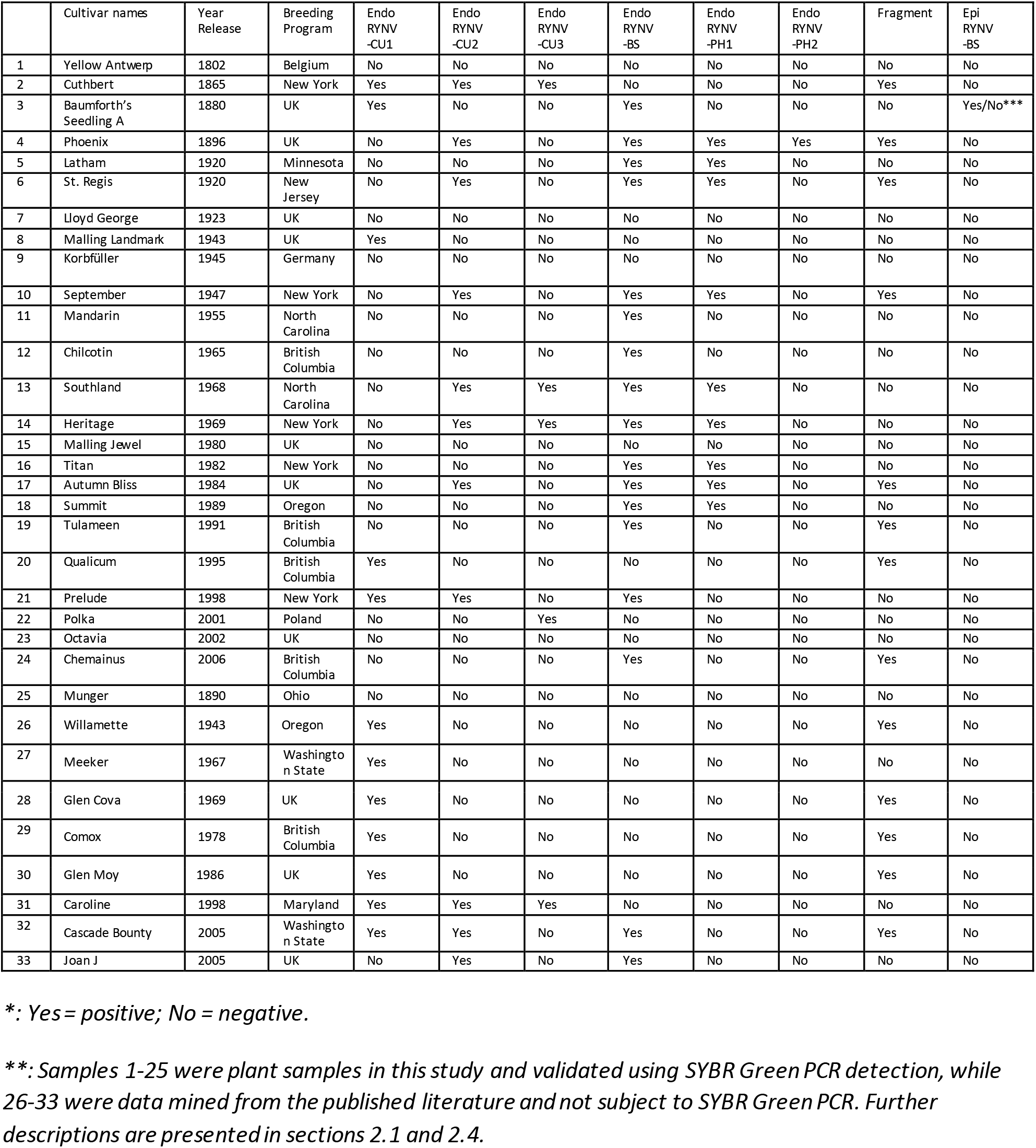
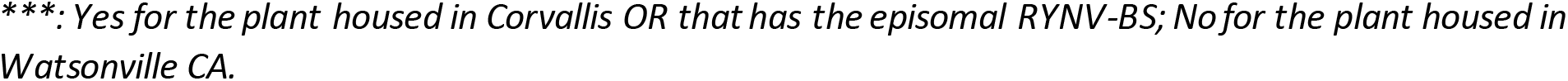
Description of the origin of the raspberry cultivars used as plants for Illumina sequencing and method development in this study, or mined from published literatures, sorted by year of release, and subsequent RYNV detection result* using analysis of whole genome sequencing data and validated by SYBR Green PCR detection.**

|  | Cultivar names | Year Release | Breeding Program | Endo RYNV -CU1 | Endo RYNV -CU2 | Endo RYNV -CU3 | Endo RYNV -BS | Endo RYNV -PH1 | Endo RYNV -PH2 | Fragment | Epi RYNV -BS |
| --- | --- | --- | --- | --- | --- | --- | --- | --- | --- | --- | --- |
| 1 | Yellow Antwerp | 1802 | Belgium | No | No | No | No | No | No | No | No |
| 2 | Cuthbert | 1865 | New York | Yes | Yes | Yes | No | No | No | Yes | No |
| 3 | Baumforth's Seedling A | 1880 | UK | Yes | No | No | Yes | No | No | No | Yes/No*** |
| 4 | Phoenix | 1896 | UK | No | Yes | No | Yes | Yes | Yes | Yes | No |
| 5 | Latham | 1920 | Minnesota | No | No | No | Yes | Yes | No | No | No |
| 6 | St. Regis | 1920 | New Jersey | No | Yes | No | Yes | Yes | No | Yes | No |
| 7 | Lloyd George | 1923 | UK | No | No | No | No | No | No | No | No |
| 8 | Malling Landmark | 1943 | UK | Yes | No | No | No | No | No | No | No |
| 9 | Korbüller | 1945 | Germany | No | No | No | No | No | No | No | No |
| 10 | September | 1947 | New York | No | Yes | No | Yes | Yes | No | Yes | No |
| 11 | Mandarin | 1955 | North Carolina | No | No | No | Yes | No | No | No | No |
| 12 | Chilcotin | 1965 | British Columbia | No | No | No | Yes | No | No | No | No |
| 13 | Southland | 1968 | North Carolina | No | Yes | Yes | Yes | Yes | No | No | No |
| 14 | Heritage | 1969 | New York | No | Yes | Yes | Yes | Yes | No | No | No |
| 15 | Malling Jewel | 1980 | UK | No | No | No | No | No | No | No | No |
| 16 | Titan | 1982 | New York | No | No | No | Yes | Yes | No | No | No |
| 17 | Autumn Bliss | 1984 | UK | No | Yes | No | Yes | Yes | No | Yes | No |
| 18 | Summit | 1989 | Oregon | No | No | No | Yes | Yes | No | No | No |
| 19 | Tulameen | 1991 | British Columbia | No | No | No | Yes | No | No | Yes | No |
| 20 | Qualicum | 1995 | British Columbia | Yes | No | No | No | No | No | Yes | No |
| 21 | Prelude | 1998 | New York | Yes | Yes | No | Yes | No | No | No | No |
| 22 | Polka | 2001 | Poland | No | No | Yes | No | No | No | No | No |
| 23 | Octavia | 2002 | UK | No | No | No | No | No | No | No | No |
| 24 | Chemainus | 2006 | British Columbia | No | No | No | Yes | No | No | Yes | No |
| 25 | Munger | 1890 | Ohio | No | No | No | No | No | No | No | No |
| 26 | Willamette | 1943 | Oregon | Yes | No | No | No | No | No | Yes | No |
| 27 | Meeker | 1967 | Washington State | Yes | No | No | No | No | No | No | No |
| 28 | Glen Cova | 1969 | UK | Yes | No | No | No | No | No | Yes | No |
| 29 | Comox | 1978 | British Columbia | Yes | No | No | No | No | No | Yes | No |
| 30 | Glen Moy | 1986 | UK | Yes | No | No | No | No | No | Yes | No |
| 31 | Caroline | 1998 | Maryland | Yes | Yes | Yes | No | No | No | No | No |
| 32 | Cascade Bounty | 2005 | Washington State | Yes | Yes | No | Yes | No | No | Yes | No |
| 33 | Joan J | 2005 | UK | No | Yes | No | Yes | No | No | No | No |

\*: Yes = positive; No = negative.
\*\*: Samples 1-25 were plant samples in this study and validated using SYBR Green PCR detection, while 26-33 were data mined from the published literature and not subject to SYBR Green PCR. Further descriptions are presented in sections 2.1 and 2.4.
\*\*\*: *Yes for the plant housed in Corvallis OR that has the episomal RYNV-BS; No for the plant housed in* *Watsonville CA.*

### 2.2. DNA purification, sequencing, and virus discovery

DNA was extracted using either the DNeasy(R) kit (Qiagen) or the method described by Poudel et al. (2013). All DNA libraries were constructed using a TruSeq DNA HT Sample Prep(R) kit and sequenced individually using paired-end (2 × 300 bp) Illumina HiSeq configuration by Novogene (Sacramento, CA). Raw Illumina reads were subject to de novo assembly using Spades (Bankevich et al., 2012). BLASTn search (Camacho et al., 2009) was performed on the output contigs with e-value=10 against published RYNV nucleotide sequences (nt) downloaded from GenBank nt database (January 16, 2021). After RYNV hits were filtered out, the remaining contigs were processed using BLASTx against a database containing all RYNV protein sequences downloaded from GenBank nr (January 16, 2021). All Illumina datasets were also submitted to VirFind (http://virfind.org, Ho and Tzanetakis, 2014) for virus detection and discovery. Bowtie2 (Langmead and Salzberg, 2012) was used for mapping raw reads to RYNV contigs for visual confirmation of the mapping assemblies with Tablet (Milne et al., 2013). BioEdit (Hall, 1999) was used to calculate sequence identity matrix, and ClustalW (Thompson et al., 1994) of the MEGA X software (Kumar et al., 2018) applied to align nucleotide and amino acid sequences. Expasy (https://web.expasy.org/translate/) was used to predict open reading frames (ORF). Conserved domain search was done using the NCBI homonymous tool (Lu et al., 2020). Breaking points of the RYNV lineages were identified by aligning raw Illumina reads with BLASTn against the assembled sequences and partially aligned reads were manually analyzed for sequence identities.

### 2.3. Electron microscopy

Tissues were homogenized in 100 mM potassium phosphate pH 7.0, 2% polyvinylpyrrolidone (MW:10,000) and 0.2% Na_2_SO_3_ at 1:20 (w:v). After a low-speed centrifugation at 10,000 g, the supernatant was used for immunosorbent electron microscopy (ISEM) according to Lockhart (1986). Briefly Formvar/carbon coated copper grids were floated on 10 μl of the capture antibodies of *s*ugarcane bacilliform virus (NanoDiagnostics, Arkansas) diluted 1:10 in 50 mM potassium phosphate buffer pH7.0. After 30 min incubation at 37°C, grids were rinsed with 50 mM potassium phosphate buffer pH7.0, and then floated on 30 μl of the sample preparations for 20-22 hrs at 4°C. The grids were then rinsed with 50 mM potassium phosphate buffer pH7.0 containing 100 μg/ml bacitracin and stained with 0.5% phosphotungstic acid pH7.0 containing 100 μg/ml bacitracin. The grids were examined using a Hitachi H-7500 transmission electron microscope (Hitachi High-Tech Corporation, Fukuoka, Japan) with an AMT Biosprint 12M-B CCD camera (Advanced Microscopy Techniques, Woburn, MA). Virus particles were measured using the camera software.

### 2.4. Data mining

Published datasets were mined for RYNV sequences (Table 1) including raw Illumina data of red raspberry cultivars ‘Caroline’, ‘Cascade Bounty’, ‘Comox’, ‘ Glen Cova’, ‘Meeker’, and ‘Willamette’ from the Diaz-Lara et al. (2020) study, ‘Glen Moy’ from Hackett et al. (2018), and the assembled ‘Joan J’ genome, obtained using PacBio and Illumina sequencing (Wight et al. 2019). These eight datasets were processed using the procedures described above.

### 2.5. Development of RYNV lineage-specific primers and validation

For each endo/epi RYNV lineage, 20 PCR primer sets were designed by processing the corresponding sequences using PrimerQuest at default parameters for ‘qPCR Intercalating Dyes’ option (Integrated DNA Technologies, IDT). The outputs were aligned with all RYNV sequences and 5-10 oligo pairs were selected with each oligo, when possible, having at least 2nt mismatches to other RYNV lineages. SYBR Green quantitative PCR was performed for each set against cultivars 1-25 (Table 1) using 20 ng plant DNA, 5 μl Maxima SYBR Green qPCR Master Mix (2X) (Thermo Scientific, catalog number K0253), 1 μM each of forward and reverse primers, and water to 10 μl. Amplification was performed on QuantStudio 6 Flex instrument (Applied Biosystems) with the amplification program consisting of 95°C for 10 min, followed by 40 cycles of 95°C, 53°C, and 72°C, for 20s each. The melting stage started at 53°C for 1m, increased by 0.05°C/s and stopped after reaching 95°C for 15s. To investigate further the possibility of integration of the epiRYNV-BS lineage, we used the most consistent assay against the epiRYNV-BS lineage developed here against a panel of 271 public and proprietary genotypes bulked into 294 DNA samples consisting of 876 plants (Supplemental table). Samples were considered positive for a lineage if there was amplification with the correct melting point. The previously published assay of Diaz-Lara et al. (2020) was also included in this validation for specificity comparison.

## 3. Results

### 3.1. A diverse array of endoRYNV is present in raspberry

The presence of RYNV DNA in commercial raspberries was investigated by whole genome sequencing and mining data of 25 and 8 cultivars, respectively (Table 1). Sequencing produced approximately 9 Gbp for each cultivar, representing ~30 X coverage of the predicted 300-Mbp raspberry genome (Wight et al., 2019). BLASTn and BLASTx steps using contigs assembled from the raspberry genomes found no evidence of RYNV DNA in cultivars ‘Korbfüller’, ‘Lloyd George’, ‘Malling Jewel’, ‘Octavia’, ‘Yellow Antwerp’, or the black raspberry cultivar ‘Munger’. The remaining 27 cultivars showed a diverse array of RYNV sequences, ranging from six partial genomic segments with duplicated and rearranged sequences, to rudimentary fragments of a few hundred base pairs. None of the newly discovered lineages possesses an intact genome. Bacilliform virus particles were consistently observed using electron microscope from ‘Baumforth’s Seedling A’ from Corvallis OR, the only sample in the study with a verified episomal form of the virus (Diaz-Lara et al., 2020) (Figure 1). The size of the virions is approximately 149 nm X 33 nm (n=102). No additional badnavirid sequences were detected in this or any other sample using VirFind (Ho and Tzanetakis, 2014) indicating that the observed particles belong to RYNV. Bacilliform particles were not detected from any of the other samples, including the ‘Baumforth’s Seedling A’ from Watsonville CA indicating that all other RYNV sequences are integrated in the raspberry genome.

**Figure 1.**
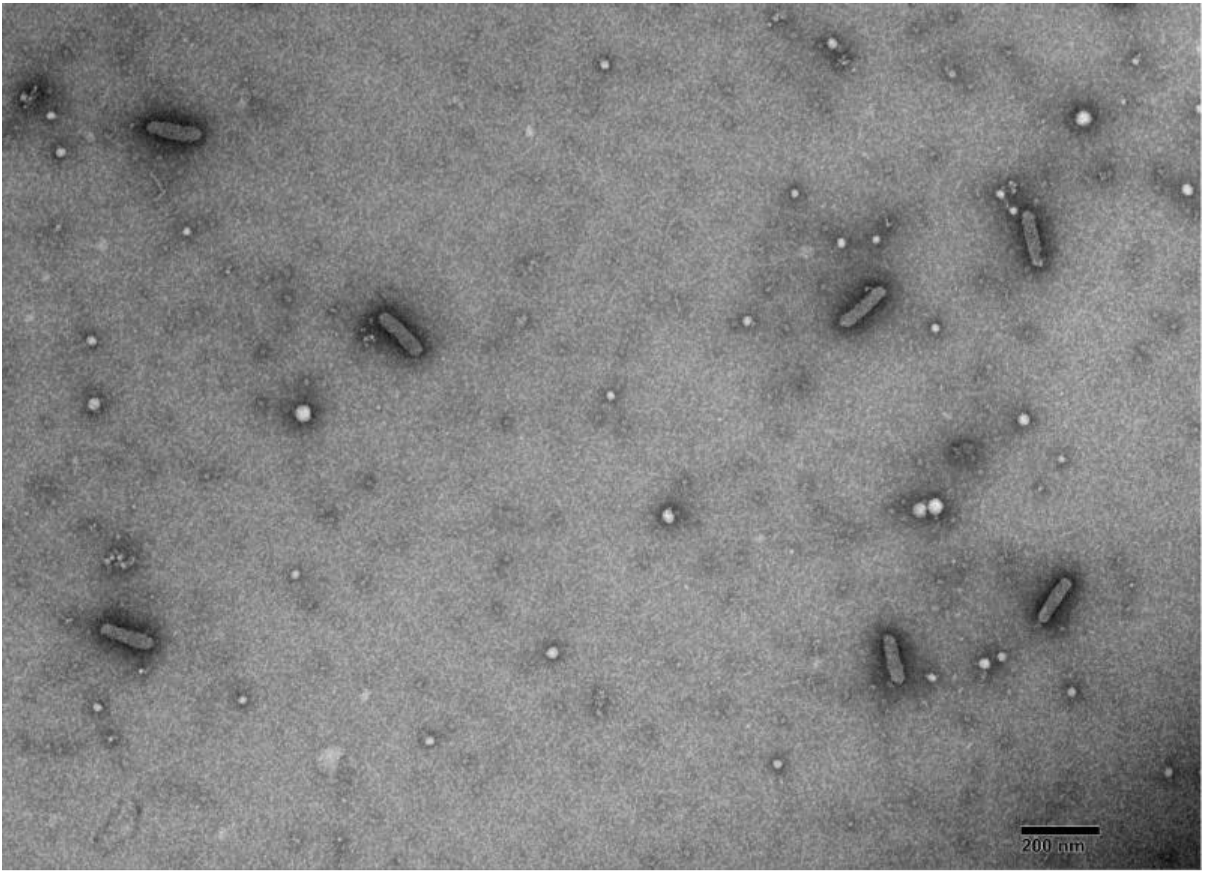
Bacilliform particles of rubus yellow net virus from ‘Baumforth’s Seedling A’ (epiRYNV-BS) captured using antibody against sugarcane bacilliform virus and stained in phosphotungstic acid.

### 3.2. Structure of main endoRYNV lineages

RYNV-derived sequences more than 4 Kbp in length were named based on the oldest cultivars they were identified in. RYNV has five conserved badnavirid domains including reverse transcriptase, ribonuclease H (RNaseH), pepsin-like aspartate protease, a zinc knuckle which is a zinc binding motif from retroviral gag proteins (nucleocapsid), and a ribosomal L25/TL5/CTC N-terminal 5S rRNA binding domain (Diaz-Lara et al., 2015). The reverse transcriptase and RNaseH (RT_RNaseH) domains were concatenated and used for sequence comparison against those of the three available genomes (GenBank accession numbers KF241951, KM078034, and MN245240). The new lineages shared >78.5% in nucleotide identities to each other and the epiRYNV-BS (Table 2).

**Table 2.** Percent identity matrix of the reverse transcriptase and RNaseH region of the six Rubus yellow net virus (RYNV) lineages discovered in this study against the three published RYNV sequences, *Gooseberry vein banding virus* (GBVBV), *Grapevine vein clearing virus* (GVCV), and *Birch leafroll associated virus* (BLRaV).

|  | endoRYNV<br>V<br>-BS | endoRYNV<br>V<br>-CU1 | endoRYNV<br>V<br>-CU2 | endoRYNV<br>V<br>-CU3 | endoRYNV<br>V<br>-PH1 | endoRYNV<br>V<br>-PH2 | epiRYNV<br>-BS<br>KM0780<br>34 | RYNV<br>-Cu<br>MN2452<br>40 | RYNV<br>-Ca<br>KF24195<br>1 | GBVBV<br>HQ8522<br>49.1 | GVCV<br>KT90747<br>8.1 |
| --- | --- | --- | --- | --- | --- | --- | --- | --- | --- | --- | --- |
| endoRYNV-CU1 | 80.5 |  |  |  |  |  |  |  |  |  |  |
| endoRYNV-CU2 | 80 | 82.5 |  |  |  |  |  |  |  |  |  |
| endoRYNV-CU3 | 79.8 | 90.3 | 81.8 |  |  |  |  |  |  |  |  |
| endoRYNV-PH1 | 80.3 | 90.3 | 81.3 | 88.5 |  |  |  |  |  |  |  |
| endoRYNV-PH2 | 81.1 | 87.1 | 89.3 | 85.2 | 85.4 |  |  |  |  |  |  |
| epiRYNV-BS<br>KM078034 | 78.5 | 81.2 | 79 | 82.1 | 84.4 | 81.3 |  |  |  |  |  |
| RYNV-Chile<br>MN245240 | 79.7 | 90.1 | 81.7 | 99.6 | 88.3 | 85 | 82 |  |  |  |  |
| RYNV-Ca<br>KF241951 | 80.5 | 100 | 82.5 | 90.3 | 90.3 | 87.1 | 81.2 | 90.1 |  |  |  |
| GBVBV<br>HQ852249.1 | 71.1 | 71 | 71.5 | 71.1 | 72.1 | 71.8 | 71 | 70.9 | 71 |  |  |
| GVCV<br>KT907478.1 | 67.7 | 68.1 | 66.7 | 67.5 | 68.4 | 66.9 | 68.5 | 67.4 | 68.1 | 68.2 |  |
| BLRaV<br>MG6864191 | 66.9 | 66.4 | 65.4 | 66.4 | 67.3 | 65.7 | 65.7 | 66.3 | 66.4 | 67.3 | 69.8 |

#### 3.2.1. endoRYNV-CU1

‘Cuthbert’ (1865 release, New York), has three endoRYNVs (namely endoRYNV-CU1, −CU2, and −CU3) and is the oldest cultivar in the study having an endoRYNV. endoRYNV-CU1 was discovered in 12 cultivars (Table 1). The 7268-nt segment has intact 5’ intergenic region (IG), ORF1, ORF2, but when compared to epiRYNV-BS, ORF3 is missing 549 bp after nt1691, corresponding to 183 amino acids (aa), at the site for the ribosomal L25/TL5/CTC N-terminal 5S rRNA binding domain, essential to the badnavirus movement (Figure 2). The four other domains are similar to the epiRYNV-BS. ORFs 4 and 6 are embedded intact within ORF3.

**Figure 2.**
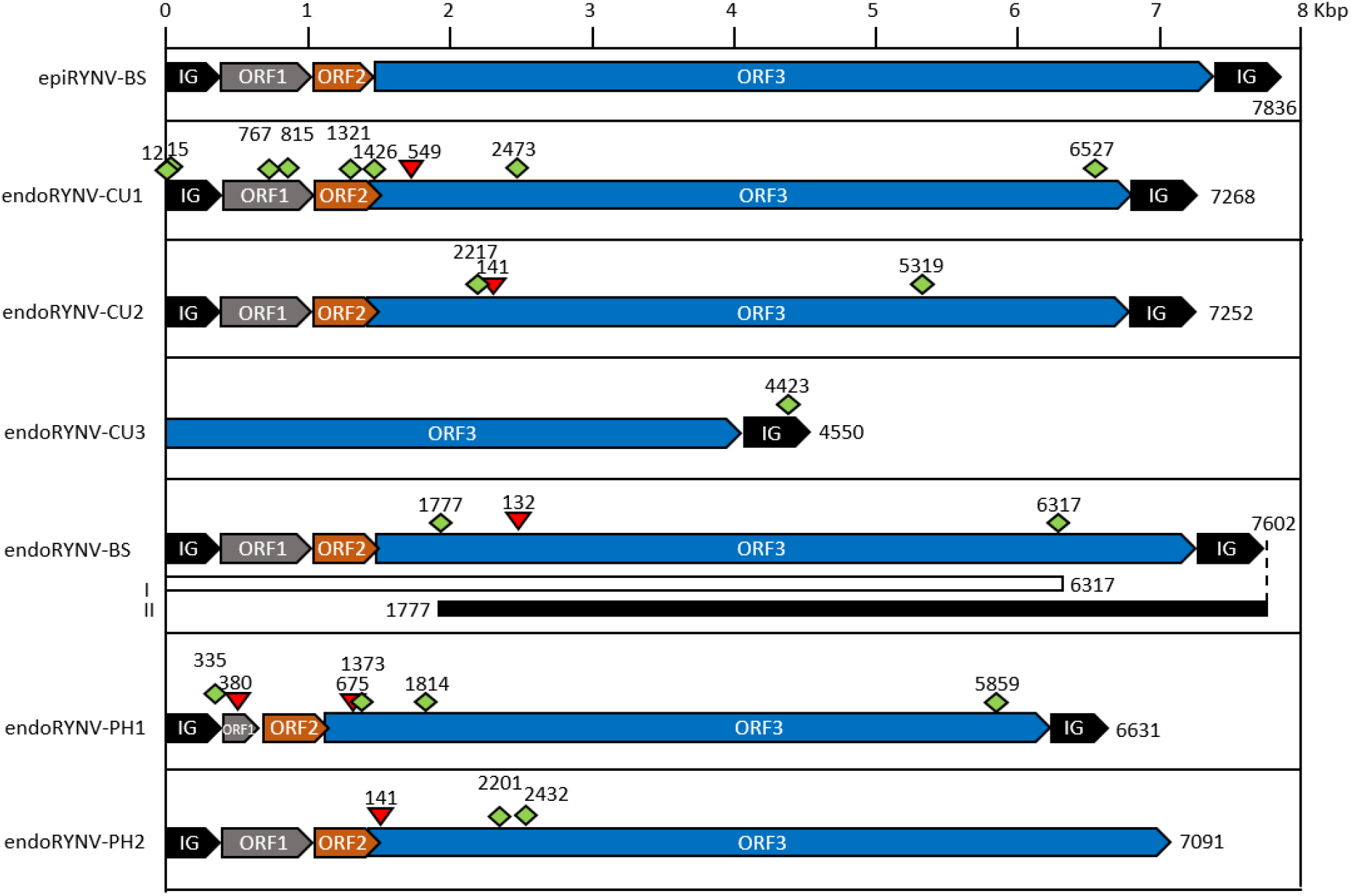
Overview of endogenous rubus yellow net virus (endoRYNV) lineage sequence structures in red raspberry. The sequences are represented with intergenic region (IG) in black; grey, orange, and blue boxes indicating ORF1, ORF2 and ORF3 of the virus, respectively. For simplicity, ORF4 and ORF6 embedded within ORF3 are not illustrated. Break points and deletions in the genomic sequence are represented in green diamond and red triangle shapes, respectively, with nucleotide location on top. The two fragments of endoRYNV-BS that were found in high-quality ‘Joan J’ genome are shown in white and black bars.

endoRYNV-CU1 RT_RNaseH region shares 100% nt identity with that of RYNV-Ca (Kalischuk et al., 2013). However, there is 12.9% nt diversity between the two lineages. endoRYNV-CU1 3’ IG is intact but very different from RYNV-Ca 3’ IG, sharing only 29% nt identity (Figure 3). The RYNV-Ca has two inverted repeats, at nt4325-4693 and nt7564-7932. The endoRYNV-CU1 integrant is highly fragmented and has breaking points at genomic nt positions 12, 15, 767, 815, 1321, 1426, 2473, and 6527 (Table 3). One plant-virus junction was detected at nt7268.

**Table 3.**
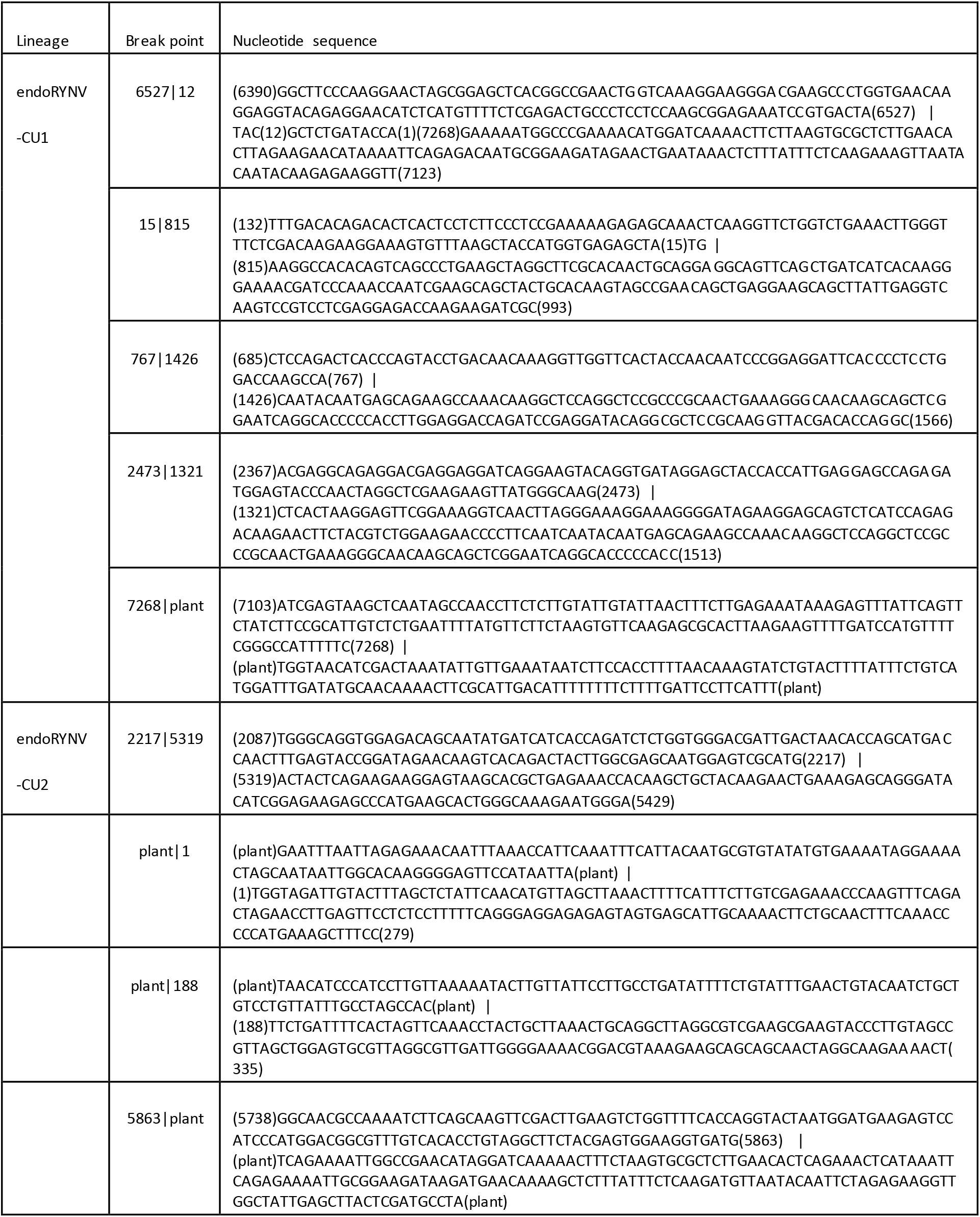
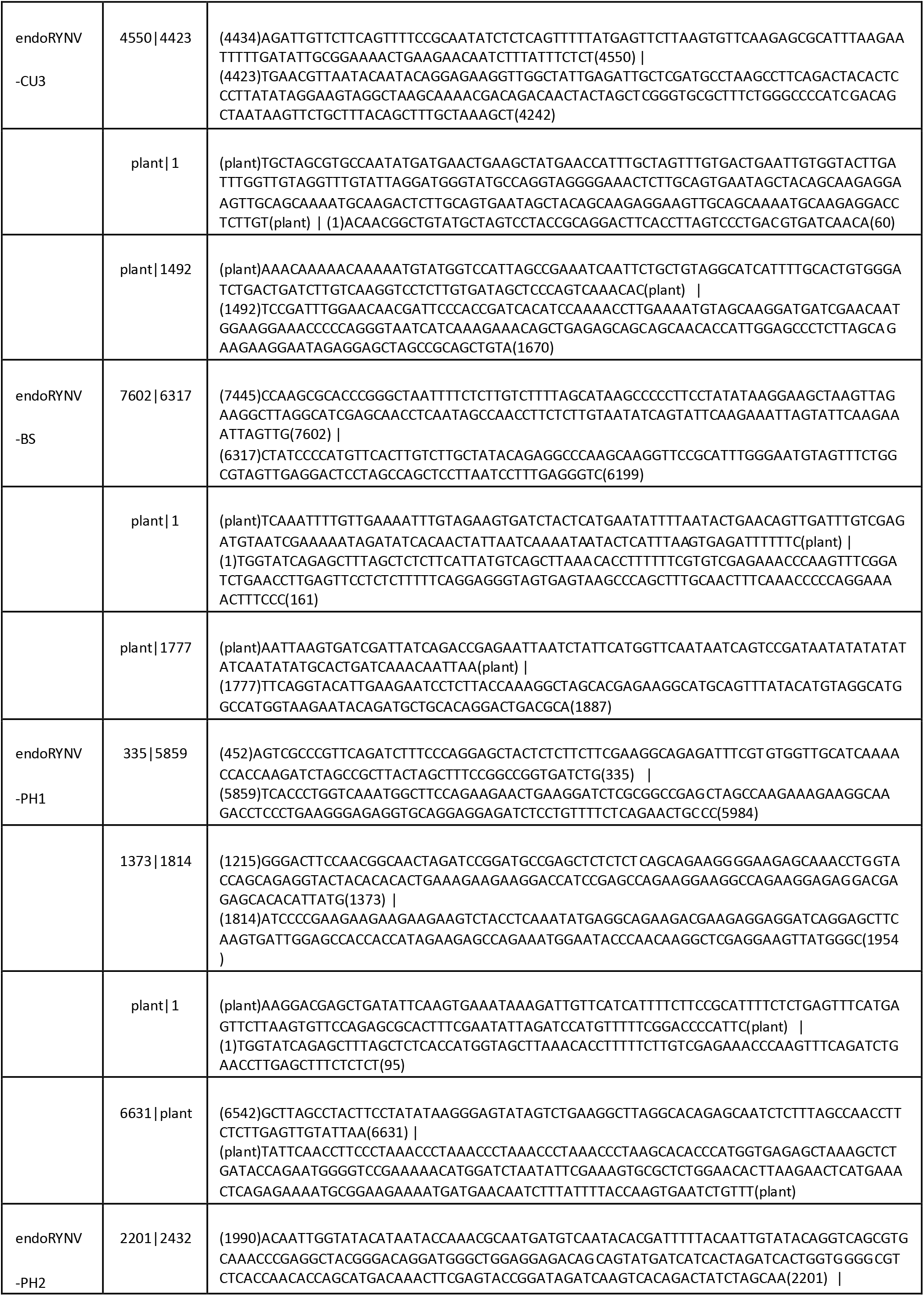
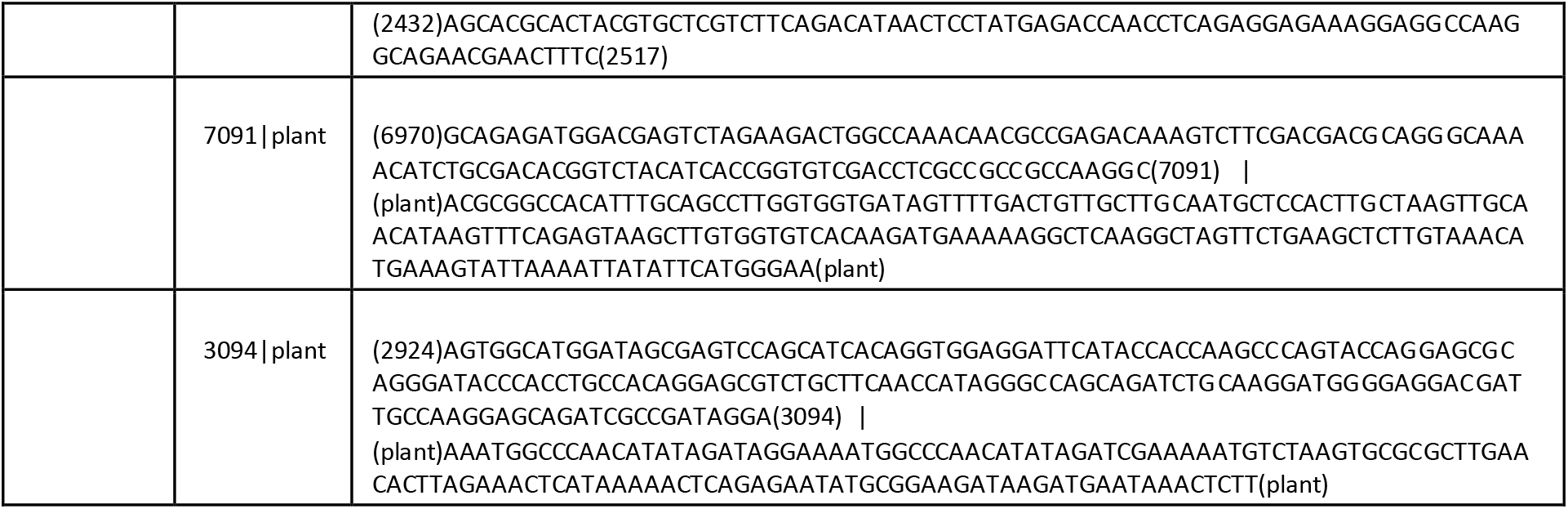
Break points in the genomic sequence, and plant-virus junctions of the endogenous RYNV lineages discovered in this study. Nucleotide positions relative to the corresponding RYNV genomic sequence are shown within parentheses. Vertical bars indicate break/junction points.

**Table 4.**
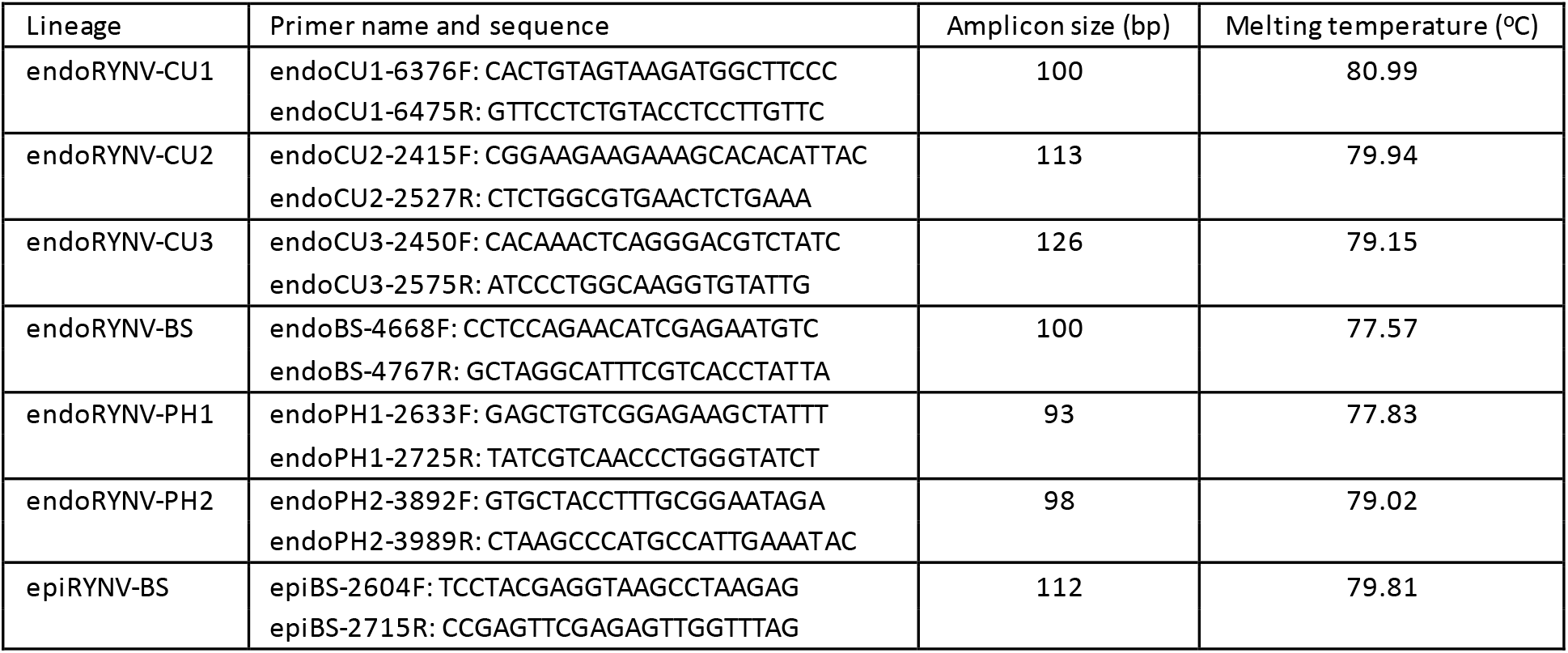
Primer pairs used for Rubus yellow net virus lineage-specific SYBR Green PCR validation.

**Figure 3.**
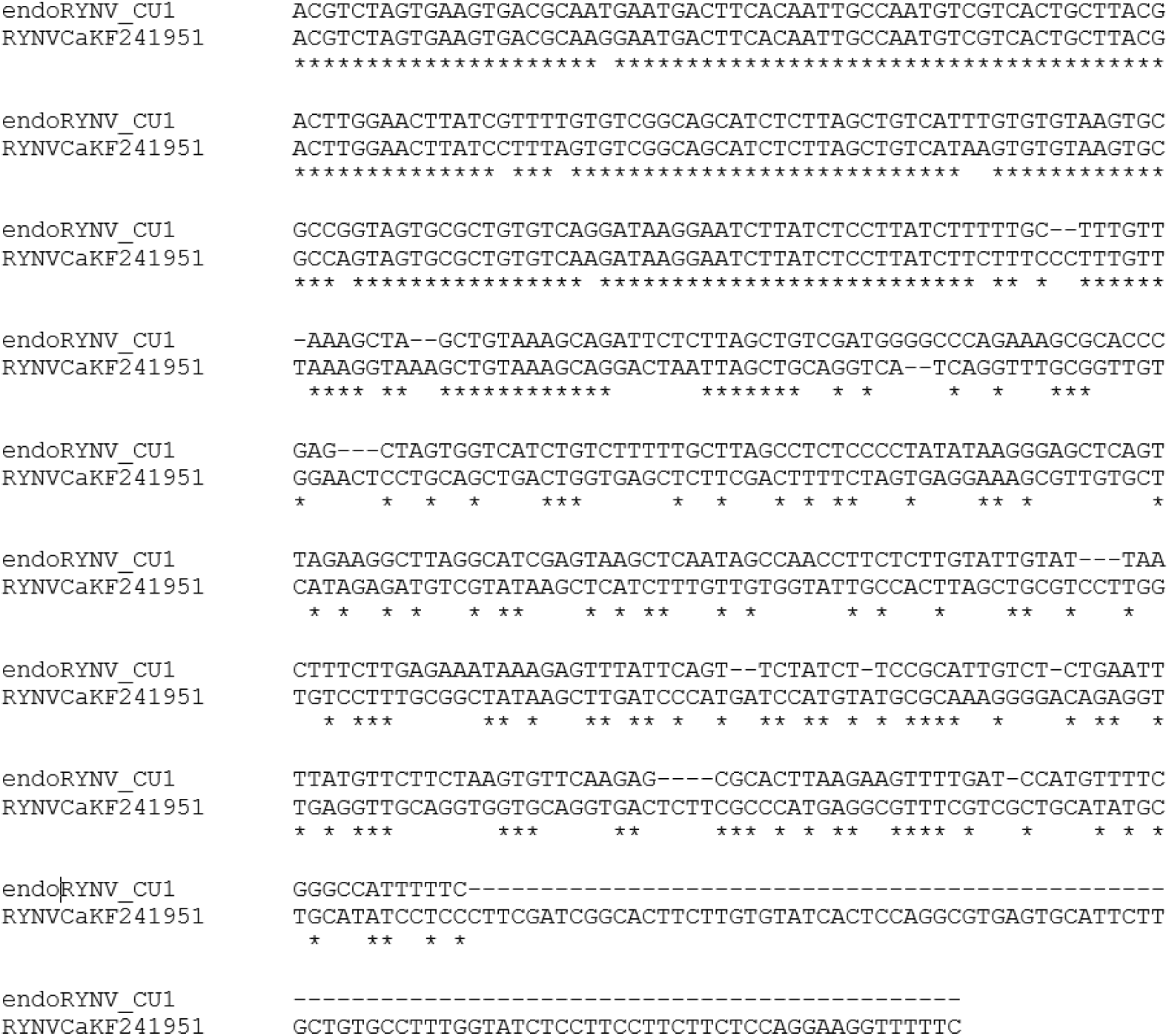
ClustalW alignment of the 3’ intergenic regions of the endogenous rubus yellow net virus (endoRYNV) lineages Cuthbert and Canada (endoRYNV-CU1 and RYNV-Ca respectively. Although the two lineages share 100% nucleotide (nt) identity at the reverse transcriptase and RNaseH domain, their 3’ IGs share 29% nt identity.

#### 3.2.2. endoRYNV-CU2

The second ‘Cuthbert’ endoRYNV (CU-2) is present in 11 cultivars. The 7252-nt sequence starts with the complete 5’ IG, ORF1 and ORF2. When aligned against epiRYNV-BS, ORF3 is lacking a 141-nt stretch after nt2455, corresponding to 47 aa. Similar to endoRYNV-CU1, it has four conserved domains similar to the epiRYNV-BS but missing the ribosomal L25/TL5/CTC N-terminal 5S rRNA binding domain as well as about 428 nt of the 3’ IG. Alike endoRYNV-CU1, its ORF4 and ORF6 are embedded intact within ORF3. The endoRYNV-CU2 RT_RNaseH region shares 79% nt identity with that of epiRYNV-BS and the integrant is fragmented at genomic nt positions 2217 and 5319, with plant-virus junctions detected at nt1, 188, and 5863.

#### 3.2.3. endoRYNV-CU3

The third ‘Cuthbert’ endoRYNV (CU-3) is present in five cultivars and is heavily truncated. It is lacking 5’ IG, ORF1 and ORF2. Its sequence starts with a truncated ORF3 and together with the 3’ IG accounting for a 4550-bp stretch. Its RT_RNaseH region shares 99.6% to RYNV-Cu from Chile (MN245240). The integrant is fragmented at nt4423 and 4550, and has plant-virus junctions at nt1 and 1492.

#### 3.2.4. endoRYNV-BS

The endoRYNV-BS was first detected in ‘Baumforth’s Seedling A’ (1880 release, UK) and is present in 17 cultivars. The lineage is 7602 bp, and has intact 5’ IG, ORF1, and ORF2. When aligned against epiRYNV-BS, ORF3 is missing a 132-nt stretch after nt2445 corresponding to 44 aa, and the 3’ IG lacks 83 bp. This lineage has all five conserved badnavirid domains. The lineage is present in the assembled genome of ‘Joan J’ as a single copy on chromosome 4. The integrant is 12,143 bp and composed of two fragments. The first is in the forward orientation and contains complete 5’ IG that forms a junction with the plant DNA at its 5’ end, followed by complete ORF1, ORF2, and part of ORF3 that is truncated at nt6317. The second follows immediately after in the reverse orientation, with a truncated 3’ IG at nt7602, then continues with ORF3 but truncated at nt1777 fusing to the plant genome. No full-length ORF3 is present in either of the fragments.

#### 3.2.5. endoRYNV-PH1

First detected in Phoenix (1896 release, UK), endoRYNV-PH1 is present in nine cultivars. The 6631nt sequence starts with the 5’ IG, followed by the ORF1 of 177 nt and missing 380 nt after nt561 when aligned against the epiRYNV-BS, before an intact ORF2. ORF3 is missing 675 nt after nt1307 as well as the ribosomal L25/TL5/CTC N-terminal 5S rRNA binding domain. It has the four other conserved domains similar to the epiRYNV-BS. The integrant is fragmented at genomic nt positions 335, 1373, 1814, and 5859, with nt1 and nt6631 connected to the plant DNA.

#### 3.2.6. endoRYNV-PH2

The last substantial integrated RYNV sequence, endoRYNV-PH2, was only found in ‘Phoenix’. The 7091-nt fragment’s 5’ end has the intact 5’ IG, ORF1 and ORF2. ORF3 misses 141 nt after nt2461 corresponding to 47 aa, and the fragment terminates at nt7091. This sequence has all conserved domains found in the epiRYNV-BS. The integrant is fragmented at nt2201 and 2432, and has two plant-virus junctions at nt3094 and 7091.

### 3.3. Validation

SYBR Green qPCR assays were able to differentiate between the epiRYNV-BS and each of the main endoRYNVs in all cultivars used in this study, either in single or multiple integration events (Figure 4, Table 1). Except the positive control ‘Baumforth’s Seedling A’ OR, the epiRYNV-BS lineage was not detected in any of the genetics used in this study (Figure 5). The epiBS-2604F/2715R assay did not have off-target melting points and amplifications, compared to the Diaz-Lara et al. (2020) assay, presently considered the better assay for RYNV detection, with off-target in 43 cases (Supplemental figure).

**Figure 4.**
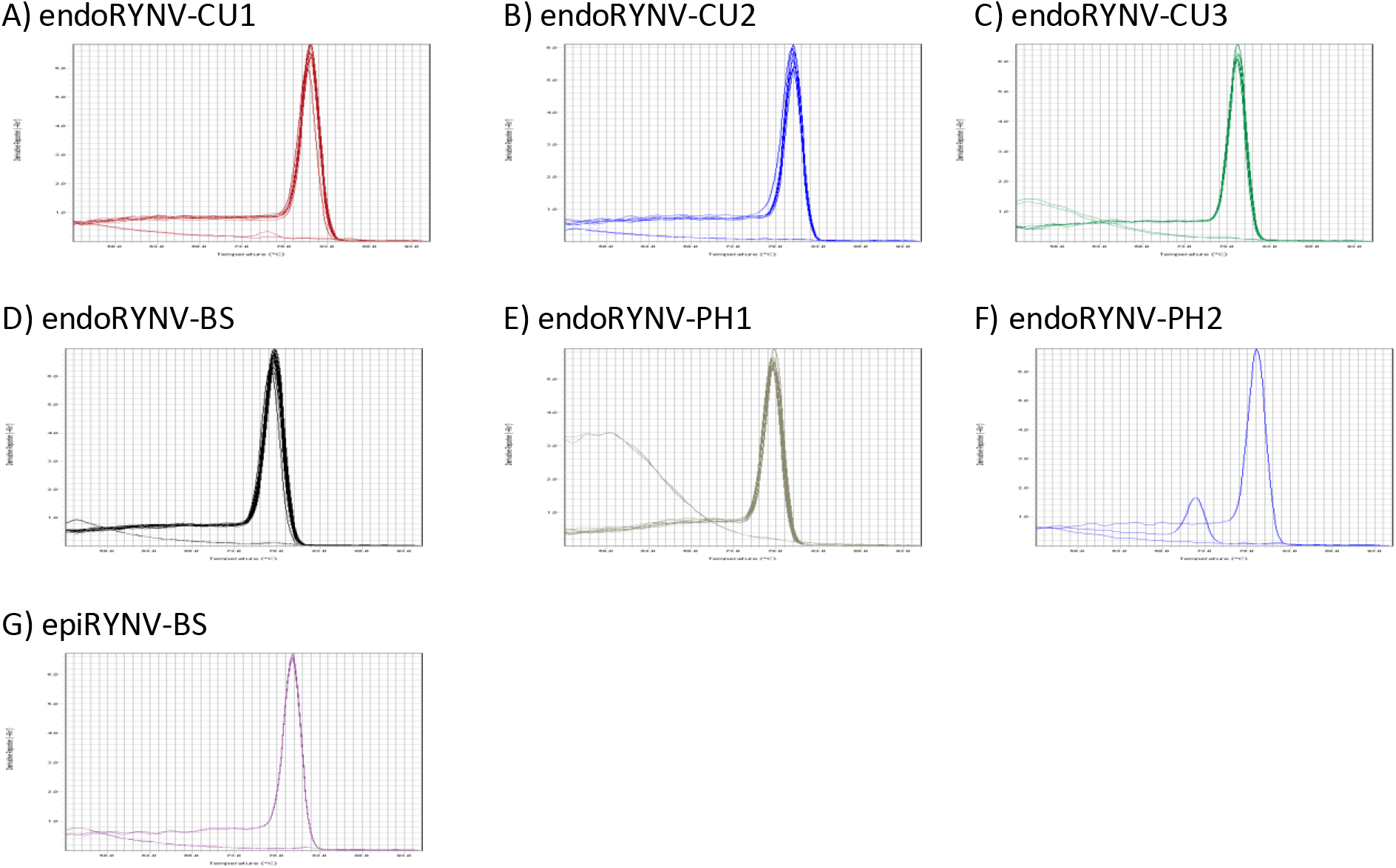
Melting curves for the lineage-specific SYBR Green PCR validation assays detecting the endogenous and episomal rubus yellow net virus (endoRYNV and epiRYNV respectively) lineages from the raspberry cultivars with corresponding endo/epiRYNVs. A) endoRYNV-CU1; B) endoRYNV-CU2; C) endoRYNV-CU3; D) endoRYNV-BS; E) endoRYNV-PH1; F) endoRYNV-PH2; G) epiRYNV-BS.

**Figure 5.**
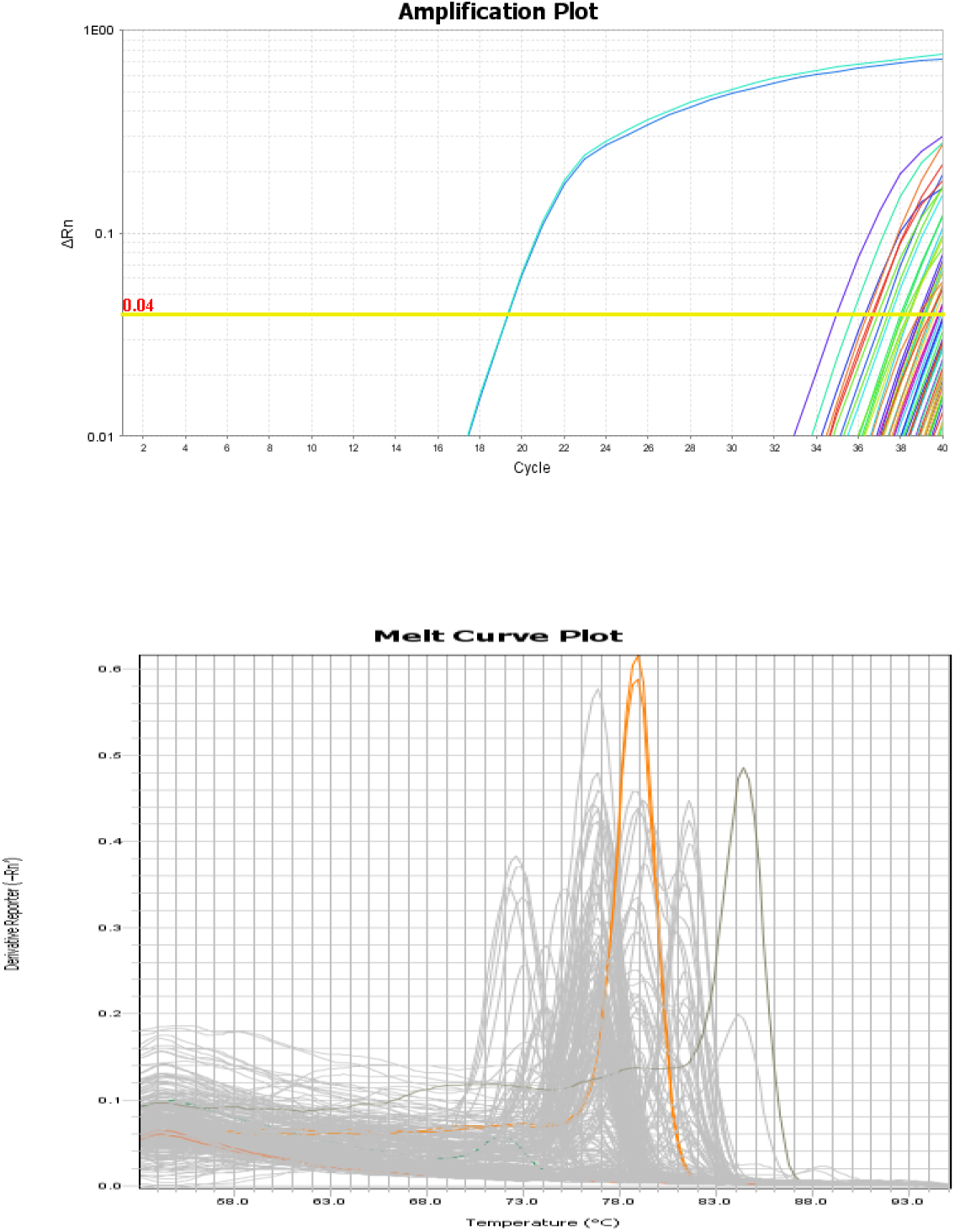
Detection of the rubus yellow net virus from ‘Baumforth’s Seedling A’ (epiRYNV-BS) lineage using the epiBS-2604F/2715R assay against a panel of 271 public and proprietary genetics. The plants were as described in 2.5. A) Amplification plot (the epiRYNV-BS positive control from Corvallis, Oregon, with two replicates were labeled in blue and light blue); B) Melting plot (the two positive control replicates were labeled in orange with unique melting point compared to all other non-specific amplifications, labeled in grey). epiRYNV-BS was detected at Ct=19, whereas non-specific amplifications initiated after Ct=35 but their melting points were different from that of the positive control.

## 4. Discussion

We analyzed the genome sequence data of commercial cultivars from around the globe released as early as 1802 and as recent as 2006. Integrated RYNV sequences were present in 27/33 cultivars (82%). The endoRYNV population could be categorized into six main lineages and other short endogenous fragments. The diversity of endoRYNV is complex with sometimes sequences having inversions, duplications, or deletions.

Rubus domestication has resulted in a reduction of genetic diversity (Haskell, 1960; Jennings, 1988), and modern cultivars are genetically similar to each other (Dale et al., 1993; Graham and McNicol, 1995). This can be seen in the case of the cultivars analyzed in this study. All breeding programs share the same endoRYNV lineages, which in turn were discovered in three cultivars commercialized in the 19^th^ century: ‘Cuthbert’ (1865), ‘Baumforth’s Seedling A’ (1880), and ‘Phoenix’ (1896). These endogenous sequences presumably became widespread as the three aforementioned cultivars were used as parents, or are in the lineages of most raspberry breeding programs worldwide.

endoRYNV-CU1 lineage is the closest isolate to the published RYNV-Ca sequence (KF241951) (Kalischuk et al., 2013). Since RYNV-Ca has two inverted repeats, misses the true 3’ IG, and hence likely is an endogenous sequence, we consider epiRYNV-BS as the sole episomal RYNV lineage known to infect *Rubus*. It is important to note that when aligned against the epiRYNV-BS sequence, all endoRYNV lineages are truncated and missing genomic DNA stretches. From this data, we hypothesize that the endoRYNVs are unable to reactivate and become episomal due to their incomplete genomes. In addition to the raspberry cultivars of this study, we sequenced the whole genome of an additional 75 proprietary red raspberry cultivars, and epiRYNV-BS was absent in all (data not shown), indicating that this lineage may be unable to integrate in the raspberry genome. We also sequenced the genomes of 100 proprietary blackberry cultivars (data not shown) but did not find any evidence of endoRYNV, suggesting that endoRYNV sequences may be limited to red raspberry.

Diagnostic tests for infectious agents are necessary so that phytosanitary agencies can protect a country’s natural resources and agriculture. However, the Rubus industry could be significantly impacted if a diagnostic test was positive for RYNV but inadvertently a no-risk endoRYNV was detected. Published PCR primers were designed to target either RYNV-Ca or epiRYNV-BS as they had been the only known RYNV lineages (Diaz-Lara et al., 2020; Jones et al., 2002; Kalischuk et al., 2008). Diaz-Lara et al. (2020) showed that primers currently used for RYNV detection could produce positive results in cultivars only harboring endoRYNV DNA, indicating the urgency to have a good diagnostic test that can clearly differentiate the two forms, similar to epiBS-2604F/2715R. This test should be developed and validated for accuracy and sensitivity against a wide range of episomal isolates.

Theoretically, endoRYNV can be removed from the red raspberry by traditional breeding. However, the effort required to remove endoRYNV DNA after multiple generations of backcrossing would be considerable, especially when desired traits must be retained. CRISPR-Cas9 could be used to remove the endoRYNVs, but for cultivars with multiple endoRYNV fragments, multiple gene-editing events will need to be done. We believe that such actions are not necessary as endoRYNV fragments could not reconstruct a full, infectious, genome.

## 6. Acknowledgement

This work could not have been accomplished without the help of various colleagues. We thank the Driscoll’s team including the raspberry breeding, molecular genetics, and greenhouse departments; and Melanie Kalischuk and Hanu Pappu for discussion of the RYNV-Ca work.

## 7. Data availability

The assembled sequences of endoRYNV-CU1, endoRYNV-CU2, endoRYNV-CU3, endoRYNV-BS, endoRYNV-PH1, and endoRYNV-PH2 were deposited on the NCBI’s GenBank under accessions XXX-XXX, and the raw Illumina reads of all cultivars sequenced in this study to the Sequence Read Archive (SRA) under accessions XXX-XXX.

## 11. Supplemental data

### >endoRYNV-CU1

TGGTATCAGAGCTTTAGCTCTCACCATGGTAGCTTAAACACTTTCCTTCTTGTCGAGAAACCCAAGTTTCAGACCAGAACCTTGAGTTTGCTCTCTTTTTCGGAGGGAAGAGGAGTGAGTGTCTGTGTCAAAACCTTGAAAGATCAAACCCCCATGAAAACTTTCCTCACGGTACCATAAGTTTTCTATCCTTCACTAGTTTGAACCTACTGCTCAAACTGCAGGCTTAGGCGTCGAAGCGAAGTACCCTTGTAGCCGTTAGCAGGAGGCGTTAGGCGTTGATTGGGGAAAACTGACGTAAAGAAGCAGCAGCAACTAGGCAAGAAACCTGACGGGTAGATCACCGGCCGGAAAGCCAGTAAGCGGCTAGATCTGGGCAGTTTTGATGCAACCTCACGAAATCTCAGCCTTCGAAGAAGAAAGCAGCTCTTGGGAAAGGTCTGAACGGGCGTATCGACAAGACTTTTTATTCAGAAATCTCAGAACGTATCCACGTTGGGAGGCAAATCAGAAAACACCCTCTCTAGACTTTCCTTGCTACCACTTCAACACAACAACCGGACCACCAGTCCACCGCACTCTCTGCAGACAAGAGAACAGTAAGGATTTACCATTTCTGGTAAACACCCTGTTCGATCTCAACATCACCGAGATCCACAATCAGGCGATTCTGGACGATAAGATCTCCAGACTCACCCAGTACCTGACAACAAAGGTTGGTTCACTACCAACAATCCCGGAGGATTCACCCCTCCTGGACCAAGCCACAATATCCTTAGATCTTCAAGCCCTCAAGGCAGATCTGAAGGAAATCAAGGCCACACAGTCAGCCCTGAAGCTAGGCTTCGCACAACTGCAGGAGGCAGTTCAGCTGATCATCACAAGGGAAAACGATCCCAAACCAATCGAAGCAGCTACTGCACAAGTAGCCGAACAGCTGAGGAAGCAGCTTATTGAGGTCAAGTCCGTCCTCGAGGAGACCAAGAAGATCGCGAGATCTCTGTCCCCCGACGGATGAACCCTAGGTGGCAGGATACTGCAACCAAGGAAACCTACCTCAAAGCCATACAAGCTACCTCATCTCTCACCTCCAACAACACAGGTCTAGGCTTCATCGAGCCACATACCTACACCGGAGGACAGCTATCTACCAACCTAGCAAAACAGAACAACACGCTCATCCAGCTGTTAGTTCAGGTGCTAGAAAAGAACCTCGACCTCGAGCAGGCAATTGTCAACCTCACAGCTCAGGTCACAAGGCTAGAAAAGACCGTCTCGGAGAAAGACACAGTCAAACTCCCAGAAAGTGTCCTCAACGACCTCACTAAGGAGTTCGGAAAGGTCAACTTAGGGAAAGGAAAGGGGATAGAAGGAGCAGTCTCATCCAGAGACAAGAACTTCTACGTCTGGAAGAACCCCTTCAATCAATACAATGAGCAGAAGCCAAACAAGGCTCCAGGCTCCGCCCGCAACTGAAAGGGCAACAAGCAGCTCGGAATCAGGCACCCCCACCTTGGAGGACCAGATCCGAGGATACAGGCGCTCCGCAAGGTTACGACACCAGGCGCAGCGAGCAATGAGAAGGACCTTCAGTAGGGACTTCAGAAACACCATAGAACGGCAACTAGACCCAGATGCCGAGCTTTCCCTCAGCAGAAGAAGGAGAGTAAACCGAGTACCAGCAGAGGTATACAGATCCGGAGAATGGGATTTACAACCCAGCAGGATCGTGGCACCACTAGCAGTCCCAACAGAAGCAAGGCTTAGCCAAAACAGGAATGGCAATATAAGCCTCAGATTCACCGACTTCCGAGATCAGAGGATCGTGGAGGAAGGAGAACCATCTGAGCCAGAAGGAAGGCCAGAAGGAGAAGATGATAGCACGCACTATGTGCTCATGTTCAACCACTCAAGGTGGGACACCTTAGGGCAACCAAGCGGGAAATATGATTACATGGTGAGGTATGATGCACCAGAACCTACCGCATGGCCAACATCCAACATCGGATGGGATGATGATAAGCCACCCAAACCGCCAAGCCCTACAAAAGGATCTTTTGAGGTAAACCTCAAAGGAGAGAAGAAACTAAAAGAGAAGGAACTCGCGGAGTTCACACCGGAGACAGATCTGGTGAGCCAGTGGTTAAGTCAGCTGTCAACATCCGCACATAATAGCGGAGCCTCAAGTTCAGACGAAGAACCAAAGTTCGACGAGGCAGAGGACGAAGACGATGTGTACAACCAGCAAACCTGGCAAAAGGAAGACAAGGAGAAAAGAGACCTGGAACTACAGGGGTGGAAACCCACCGGGAGACCAGGAATCTACGAGATGATCCCCGAAGAAGAAGAAGAAATCTACCTCAGGTACGAGGCAGAGGACGAGGAGGATCAGGAAGTACAGGTGATAGGAGCTACCACCATTGAGGAGCCAGAGATGGAGTACCCAACTAGGCTCGAAGAAGTTATGGGCAAGCTCAAAAACGTGAGCATGGAAAAACTGTTCCCAGTAAGCGGAATGGACAGCGAATCCAGCATCACAGGTGGAGGATTCATCCCACCAAGCCCAGTGCCAGGAGCACAAGGGTACCCACCAGCAACTGGAGCATCCGCGTCCACCATTGGACCAGCAGACATGCAAGGATGGGGAGGACGGCTACCTCGGAGCAGGTCGCCTATAGGCTATGGCAGACCCCAACAACCGTGGTCACTGCCCTCAGCACAGTCTGATAACGGCTGCATGCTAGTCCTTCCACAGGACTTCACCCTAGTCCCCGACGTAATCAACAGATGGGAATCCATCACAGTCAACCTCATCAACAAGATGATGTTTGATTCCCTACAGGACAAGGCGGACTACGTAGAAAACCTCCTTGGAGAAAGAGAAAAGGAGACATGGATGACATGGAGAATGCAGTACGAGGAAGAGTACAGGCAACTCCTCACCATGAGCGGAGACGTAAGGAACCTTACTGCCGCAGTCAAAAGGGTCTTTGGAGTACACGACCCGCACACAGGATCAGTACACATCCAGAATCAAGCGTACGCAGAGCTGGAACGCCTCTACTGCAAAAGAACGGATGATGTGATCCCCTTCCTCTACGACTACTACCAGTTAGCAGCCAAGTCAGGAAGGATGTGGCTCGGACCTGAGCTATCTGAGAAGCTGTTCAGAAAGCTTCCACCGGAGATAGGCCCAACAATAGAGCAGGCCTATAAAGACAGGTATCCAGGCCTCACGATTGGAGTTTTGGCAAGGGCCAATTTCATCCTGGAATATCTACAAAACGTCTGCAAGCAAGCAGCGTTGCAAAGGTCCCTAAAAAGCCTGAGCTTCTGCAGAAACATGCCAGTACCAGGGTACTACGAGAAGAAGCAATACGGCATCAGAAAGGCTAAAACCTATAAAGGAAAGCCTCACCCTACCCACGTGAAAGTCATCAAAAACAAGTACAAGCACACATCTGGGAAGAAGTGCAAATGCTACTTATGTGGGATAGAAGGCCATTACGCCAGGGAATGCCCAAAGAAAGTGGTGAAGCCACAAAGAGCGGCATACTTCAATGGCATGGGACTAGACGACAACTGGGATGTCGTGTCCGTCGAGCCCGGAGAATCAGATGACGATGAAATCTGTAGCATCTCCGAGGGAGAAAACGCTGGAGGAATGCATGAGCTTATGGCATTCAAGACTCAACTCCCATACCCAGTGGAGTACGAAGCCAGCACACCACAGTTCCTGATGCCATGGACACAAGTAACAGTGGAAAGAAGCGAAAAACCTTCCTGGAGAAGAAGGAAGGAAATCCCGAAGGCACAACAGGATTGTACTCACACCTGGAGTGACACACAAGAAGTGCCTATCGAGGGAAGGATATGCAGCATATGCAGTGATGAAACCCCTCATGGGCGAAGGATCACCTGCACCACCTGCAGCCTTAACCTCTGTCCGCTTTGCGCTTACATGGATCATGGGATCAAGCTTATAGCCGCAAAGGACACCAAGGACGCAGCTAAGTGGCAATACCACAACAAAGATGAGCTTA TACGACATCTCTATGAGCACAACGCTTTCCTCACTAGAAAAGTCGAAGAGCTCACCAGTCAGCTGCAGGAGTTCCACAACCGCAAACCTGATGACCTGATCAGCTTAGCGGATGACTTGGAGGACGTGTCCATTCTGGACAACGCCTCAAAAAGGGGGAAGGAGAAGGAATCTTTCCAATTCGGAACAACGATTCCCATCGACCACATCCAAAACTTGGAAAACGTGGCAAGGATCATCGAGCAATGGAAGGATACCCCCAAGGTAATCATCAAAGAAACAGCTGAAAGCAGCAACAACACCATCGGAGCCCTCTTAGCAGAAGAAGGAATAGAGGAGCTAGCCGCAGCTGTAGACACGGCATACACAGAAATGCCAAAAGGAGGATTGAACAAGCTCTACAACACCATTGTTGAGTTTGTAATACCCCAGGAAAAGGGGGCACCCACCAGGTTCAGGGTAAGAGCTGTAATAGACACAGGATGCACCTGTACATGTATCAACAGCAAGAAAGTCCCCAAAGAAGCCCTGGAGGAAGCGAAGTACCAGATGAACTTCGCAGGAGTAAATTCCACTGGAGAAACGAAGCTTAAAATGAAGAACGGTAAGATGATCGTGTCTGGAAGCGATTTCTATACACCGTACATTGCAGCCTTCCCAATGGAACTACCAGACGTAGACATGCTCATCGGCTGCAACTTCTTGCGAGCCATGAAGGGAGGAGTCAGGCTTGAAGGTACTGAAGTGACGATCTACAAGAAAGTCACCACAATCCAAACAACCCTGGAGCCCCAAAAGATATCTCTGCTCCGCGCAGAAGCAGAAGTCGGAGAAGAGATCGAGCGTATGTACTACGCAAATGACTACTCTGAAGAAGGAGTCAGTCGCCTGAGAAACCACAAACTGCTGCAGGAACTAAAAGAACAAGGCTACATAGGCGAAGAGCCAATGAAGCACTGGGCGAAAAACGGGATCAAGTGTAAGCTTGACATCAAGAACCCAGACATAGTAATCAGCAGTAAACCCCCGGATGCTGTCTCAAAGGAGACGAAGGCACAATACCAGCGGCACATTGACGCTCTCCTGAAGATCAAAGTAATCCAGCCAAGCAAGAGCAAGCACAGAACCGCAGCCTTCATCACAAACTCGGGCACAACCGTTGACCCGATCACAAAGAAAGAAATCCGAGGAAAAGAAAGGATGGTGTTCGACTACAGAAGTCTGAACGACAACACCCACAAAGACCAGTATACTTTGCCTGGGATCAACACCATCATATCGGCAATCGGCAATGCGAAGATCTTCAGCAAATTTGATCTGAAGTCTGGATTCCACCAAGTATTGATGGACGAAGAATCCATCCCGTGGACCGCATTTGTCACACCAGTAGGGTTCTACGAGTGGAAGGTAATGCCTTTCGGACTCGCAAACGCTCCGGCCGTCTTCCAGAGAAAGATGGACCAGTGTTTTGCAGGAACCTCAGAGTTCATAGCCGTCTACATCGACGATATCCTGGTCTTCAGCAAGACCTTGAAGGAGCACGAAAAGCACCTGAGCATCATGCTTGGGATATGTCGAGACAACGGCCTGGTTTTGTCACCAAGCAAGATGAAGTTAGCAGCAACCGAGATCGACTTCTTGGGAGCCACCATTGGTGACGGAAAGATTAAACTCCAGCCTCACATAATCAAGAAGATAGCTGAGGTGGACGATGAATCTCTAAAAACCCTCAAAGGGTTGAGAAGTTGGTTGGGAGTTCTCAACTATGCCAGGAACTACATCCCGAAGTGCGGAACACTCCTAGGCCCACTATACAGCAAGACCAGTGAGCATGGAGACAGAAGGTGGCATGCTTCGGATTGGGCCTTAGTAAAGAAGATCAAGAGCCTGGTCCAAAATCTCCCAGACCTCAAACTGCCCAGTGAGGAGGCCTATATGATCATCGAGACAGATGGTTGTATGGAAGGATGGGGCGGAGTCTGTAAGTGGAAGCCCAACAAAGCAGACTCAGCTGGCAAGGAAGAAATCTGCGCTTACGCAAGCGGTAAGTTCCCAACGGTGAAATCTACCATTGACGCAGAAATCTTCGCGGTAATGGAGTCCTTAGAAAAATTCAAAATTTTCTACATGAATAAGGACGAGATCACCATCAGAACCGATTGCCACGCCATCATCACCTTTTACGAAAAGTTAAACGCCAAGAAACCTTCTCGGGTAAGGTGGTTAGCTTTTTGTGATTATATAACAAACTCAGGGGTGAAGATGAAGTTCGAACACATCAAAGGCAAAGATAATCAGCTCGCTGACAATCTTAGTCGCCTTACCCAACTCATCACTGTAGTAAGATGGCTTCCCAAGGAACTAGCGGAGCTCACGGCCGAACTGGTCAAAGGAAGGGACGAAGCCCTGGTGAACAAGGAGGTACAGAGGAACATCTCATGTTTTCTCGAGACTGCCCTCCTCCAAGCGGAGAAATCCGTGACTACTCGCCCATCAGAGCCGCACCATGTACTATGGCGGAGATGGACGAATCCCGAAGAGTGGCCATGCAGCGAAGAATCAAGGTCTTCGACGATCTTGCACAAAACATCAGCGACGCCGTATACATCACAGGCATCGACCTCGCCGCCGCCAAGGCACGGGCAACCAGAGACAACTGGTACAATGACGTCACCCCAGCATTGGAAGAACGAGCAGCTGCAGCATGGAGACTCATGGCGGCCTACTCAGACTTCGCCACGTGGAAGGACGTAAACGTCTAGTGAAGTGACGCAATGAATGACTTCACAATTGCCAATGTCGTCACTGCTTACGACTTGGAACTTATCGTTTTGTGTCGGCAGCATCTCTTAGCTGTCATTTGTGTGTAAGTGCGCCGGTAGTGCGCTGTGTCAGGATAAGGAATCTTATCTCCTTATCTTTTTGCTTTGTTAAAGCTAGCTGTAAAGCAGATTCTCTTAGCTGTCGATGGGGCCCAGAAAGCGCACCCGAGCTAGTGGTCATCTGTCTTTTTGCTTAGCCTCTCCCCTATATAAGGGAGCTCAGTTAGAAGGCTTAGGCATCGAGTAAGCTCAATAGCCAACCTTCTCTTGTATTGTATTAACTTTCTTGAGAAATAAAGAGTTTATTCAGTTCTATCTTCCGCATTGTCTCTGAATTTTATGTTCTTCTAAGTGTTCAAGAGCGCACTTAAGAAGTTTTGATCCATGTTTTCGGGCCATTTTTC

### >endoRYNV-CU2

TGGTAGATTGTACTTTAGCTCTATTCAACATGTTAGCTTAAACTTTTCATTTCTTGTCGAGAAACCCAAGTTTCAGACTAGAACCTTGAGTTCCTCTCCTTTTTCAGGGAGGAGAGAGTAGTGAGCATTGCAAAACTTCTGCAACTTTCAAACCCCCATGAAAGCTTTCCTCACGGTACTATAAGCTTTCTGATTTTCACTAGTTCAAACCTACTGCTTAAACTGCAGGCTTAGGCGTCGAAGCGAAGTACCCTTGTAGCCGTTAGCTGGAGTGCGTTAGGCGTTGATTGGGGAAAACGGACGTAAAGAAGCAGCAGCAACTAGGCAAGAAAACTGCAAATCAGATCACCGGCCGGAAAGTCAGTAAGCGGCTAGATCGGGGCAGAAGCGGATGCAACCTCACGAAATCTCATCCTTTGAAGAAGAGAGCAACTCTTGGGAAAGGTCTGAACGGGCGTATCGACAAGACTTCCTATTCAGAAATCTTAGATCCTATCCGAGGTACGAAGCAAACCAGAAATCACCTTCCTGTGATTTCCCTTGTTATCACTTCAACACAACCACTGGACCTCCAGTCCACCGCACGATCTGCAAGCAGAAGAACAGTGAGGATTTACCCTACCTGGTAAATACACTGTTTGATCTCAACATCACTGAGATCCACAACCAGGCAGTCCTAGACGACAAAGTCTCGAGACTCACACAGTACCTGGTCAAGAAAGTCGGAACGCTACCGACAATCCCGGAGGACTCACCCCTCCTGGACCAAGGTTCCATAAACCTAGATCTGCAGGCCCTAAAAGCAGATCTGAAAGAAGTCAAGGCAACCCAGTCCGCACTGAGGTTAGGCTTTGAACAGCTAAGAGAAGCTGTCCAGCTAATCATTGCCCGCGAAAACGATCCGAAGCCCATCGAAGC TTCTACGGCACTCGTAGCGGAGCAGCTGAGGAAACAACTGATAGAGGTGAAATCTGTCCTCGAGGAGACAAAGAAGATCGCCAGATCTCTCTCCCCTGACGGATGAACCCAAAGTGGCAGGAGACCGCCGCAAAAGAAACCTATCAGAAGGCTATCCAGGCAACCTCCTCGCTCACATCAAACAACACCGGTCTAGGGTTCATAGAACCTCATACTTACACCGGAGGACAACTGTCCACAAACCTAGCAAAGCAGAACAACACGCTCATACAGCTGTTGGTCCAAGTGCTAGAAAAGAATCTGGACCTTGAACAGGCGATAGCAAACCTATCAGCACAGGTCACCAGACTCGAGAAGACCGTCGCAGAGAAGGACACGGTCAAACTCCCGGAAAGCGTCCTGAACGACCTCACCAAAGAATTCGGGAAGGTCAATCTCGGGAAAGGAAAGGGTTTAGAAGGTATTGTCTCTTCCAAAGACAAAAACTTCTACGTTTGGAAGAACCCTTTCAACCAGTACAATGAGCAGAAGCCAAACAAGGCTCCAGGCTCCGCCCGCAACTGAAAGGGCAACAAGCAGCTCGGACTCAGGAACCCCCACCCTGGAGGACCAGATCCGAGGATATAGACGCTCTGCAAGGTTAAGACACCAAGCACAGCGAGCAGTTCGAAGGACCTTCAGCAGGGACTTCAGGAATACTATAGAAAGGCAACTGGACCCAGATGCCGAACTCTCCCTTAGTAGGAGAAGAAGAGCGAATCTAGTACCAGCAGAGGTACTCTATGCTCACAACGGTTCGGAACCAGCGAATCGTGTGTACGAGCATTACAGTGAGCTCGGGGCCCACATAGTCGACAGGCAACAAGACTTCCGGTATATAGAGGAAGCCTCCTACCAAAGGCTAGTCAGGGAAGGCATGCAGTTCATTCATGTCGGGATGGCCATGGTCAGGATCCAGATGTTGCATAGAACGGATGCGGGGATATCCGCATTAGTGGTGTTCAGAGACACCAGGTGGAGTGATGACAGGCAAGTCATCGGGAGCATGTCCGTGGACATGACAAGGGGAGCACAGTTGGTGTATATAATACCCAACGCAATGATGTCAGTACACGATTTCTACAATCGTATACAAGTAAGTGTGCAGACCCGAGGCTACGGGACAGGATGGGCAGGTGGAGACAGCAATATGATCATCACCAGATCTCTGGTGGGACGATTGACTAACACCAGCATGACCAACTTTGAGTACCGGATAGAACAAGTCACAGACTACTTGGCGAGCAATGGAGTCGCATGCATACCCGGACAGAAGTGGGACGTAGCCAATCGATCCGGAGAATGGGAACTTCAACCCAGCAGGATCACAGCGCCAGTCATGGCACCTACTGAGGCAAGGTTGAGCCAGAACAGAAATGGCAGCATAAGCCTCAGATTCTCAGATTTCCGAGATCAGAGAATCACAGAAGAAAGGCCAGCTGAAGAAGAGGGCAGGCCGGAAGAAGAAAGCACACATTACGTGCTCATGTTCCGTCACAGCTCCTACGAGACCAATCTCAGAGGAGAAAGGAGGCCAAGGCAGAGTGAGCTTTCAGAGTTCACGCCAGAGACAGATCTGGTAAGCCAGTGGCTGAGCCAACTATCAGCCTCAGCACACAACAGTGGAGCATCTAGTTCAGAAGACGAACCGCCCAGGTTTGATGACTCTGAAGAAGATAGTGACAACACCTATAATGAGAAGACCTGGCAAAGAGAAGACCAGGAAAGGCGAGATCTGGAGCTGCAAGGATGGAAGAAGACCAGCAGACCAGGAATCTATGAGCTGATCCCAGAGGAAGAGGAAGAAATCTACCTCAGGTATGAAGATGAAGAGGATCAGACAACAACACAGGTAATCGGGGCAACAACCATGGAGGAACCAGAGATGGAGTACCCTACCAGGTTGGAGGAAGTGATGGGAAGACTCAAAAACGTTAGCATGGAGAAGCTGTTCCCAGTTAGCGGCATGGATAGCGAGTCAAGCATCACAGGGGGAGGCTTTATCCCACCAAGCCCAGTACCAGGAGCGCAGGGTTACCCACCTGCAACAGGAGCATCCTTTGGCTCGACAATTGGGCCAGCAGATCTACAAGGATGGGGAGGTAGGCTGCCACGCAGCAGATCACCACTTGGATATGGCCGCCCACAGCAGCCATGGTCATTACCATCCGCCCAATCTGAGAACGGATGTATGCTAGTCCTCCCACAGGACTTCACTCTAGTCCCAGATGTAATAAACAGATGGGAATCCGTAACCGTCAACCTTATCAACAAGATGATGTTCGACTCCTTACAAGATAAGGCGGATTACGTGGAAAATCTCCTAGGAGAACGGGAGAAGGAGACATGGATGACCTGGAGGATGCAATATGAAGAAGAGTACAAGCAGCTCCTCACCATGGCAGGAGACGTGAGAAATCTCACGGCAGCAGTCAAGAGGGTCTTTGGAGTGCATGACCCCCATACAGGATCAGTCCACATACAAAACCAGGCATATGCAGAACTGGAGAGGCTCTACTGCAAGAGAACGGACGACGTGATCCCTTTCCTATATGACTACTACCAACTGGCGGCGAAATCTGGAAGAATGTGGCTAGGGCCAGAATTATCAGAAAAGCTGTTCAGAAAGCTGCCACCAGAGATAGGCCCAACGATTGAGCAGGCCTATAAAGACAGGTATCCAGGGCTGACAATTGGAGTCTTGGCGCGAGCCAACTTCATACTGGAGTACCTACAGAATGTCTGCAAGCAGGCAGCGCTACAGAGGTCTTTGAAGAGCCTCAGCTTTTGTAGAACCATGCCGGTGCCAGGGTATTATGAGAAGAAGCAATACGGCATCAGGAAGGCAAAGACCTACAAAGGGAAGCCACACCCCACACATGTAAAGGTAATCAAGAACAAGTATAAACACTCCGCAGGGAGGAAATGCAAATGTTACCTTTGCGGGATAGAGGGTCACTACGCCAGGGAATGCCCAAAGCAAGTGGTTAAGCCACAGAGAGCAGCATTCTTTAATGGCATGGGCCTTGATGATAACTGGGACGTCGTGTCCGTGGAGCCAGGAGAGACAGATGATGATGAGATCTGCAGCATCTCAGAAGGCGAGGGCACCGGGGGGATGAACGAGCTACTTGCATTCAAGACACAGCTCCCCTACCCAGTAGAATATGAGGCGAGCTCATCTCAGCAGTTCCTGCCATGGATTCAGGTAACTGTACAAAAGAGTGAGAAACCCTCATGGAGAAGAAGGAAAGAGATCCAACCGGCACAACAAGAATGCGCTCACCTCTGGAGCGATACACAAGAAGTGCCTATAGAAGGAAGAATTTGCAGCATCTGCAGTGACGAAACCCCTCTGGGAAGAAGGATCACCTGCACTGCCTGCAACCTAAATCTGTGTCCTCTTTGCGCATACATGGACCATGGTATCCAACTGATAGCCGCAAAGGACACCAGAGATGCAGCAAAATGGCAGTACCACAACAAGGATGAGCTAATCCGGCATCTCTATGAGCATAATGCTTTCTTAACCAGAAAGGTGGAGGAGCTGACAAGCCAGCTGCAACAATGGCAAAACCGCAAACCAGAAGACCTTATCAGTCTTGCGGATGAGCCAGAGGACGTGTCCATCCTGGACAACGCCTCAAAAAACCGGGGGAAGGAGAAGGATTCTTTCCAATTTGGAACAACAATCCCAGTCGATCATATACAAAACATCGAGAATGTTGGCAGAATGATCGAACAATGGAAGAACACCCCCAATGTCACGGTCAAAGAAGTGGCAGAATCAAGCAGCAACACCATAGGAGCACTCCTTGCAGAAGAAGGAATTGAAGAGCTCGCGGCTGCAGTAGATACGGCATACACTGAAATGCCGAAAGGAGGGTTGAACAAACTCTACAACACCATCGTAGAATTTGTGATACCACAGGAAAAGGGGGCACCCTCTAGATTCCGGGTCAGAGCTGTCATTGACACAGGATGCACTTGTACCTGCATCAACAGCAGGAAGGTCCCGAAGGAAGCTCTCGAGGAAGCGAAGTTTCAGATGAATTTCGCCGGGGTAAACTCCACGGGAGAAACAAAGCTGAAAATGAAGAATGGCAAAATGATAGTATCAGGGAGTGATTTCTATACGCCATACATCGCGGCTTTCCCCATGGAACTACCAGATGTGGACATGCTCATTGGCTGTAACTTCTTACGAGCCATGAAAGGAGGAGTCAGACTTGAAGGAACAGAAGTAACCATCTACAAGAAGGTTACCACAATCCAGACGACTCTAGAGCCACAGAAGATCTCTCTACTCCGTGCAGAGGCA GAAGCAGGGGAAGAAATTGAGAGACTCTACTATGCCAATGACTACTCAGAAGAAGGAGTAAGCAGGCTGAGAAACCACAAGCTGCTACAAGAACTGAGAGAGCAGGGATACATCGGAGAAGAGCCCATGAAGCACTGGGCAAAGAATGGGATCAAGTGCAAACTTGATATCAAAAATCCCGATATAGTTATCAACAGCAAGCCTCCGGATGCAGTGTCTAAAGAGACGAAGGCGCAGTACCAAAGGCACATCGATGCTTTACTGAAGATCAAGGTGATTCAACCCAGTAAAAGTAAGCACCGAACAGCAGCCTTCATCACCAACTCGGGTACCAGCATAGACCCGATAACAAAGAAGGAGATCCGAGGAAAGGAGAGGATGGTCTTCGACTACAGGAGTCTGAATACCCACAAGGATCAGTACACCTTGCCTGGGATAAATACCATCATCTCTGCAATTGGCAACGCCAAAATCTTCAGCAAGTTCGACTTGAAGTCTGGTTTTCACCAGGTACTAATGGATGAAGAGTCCATCCCATGGACGACGTTTGTCACACCTGTAGGCTTCTACGAGTGGAAGGTGATGCCTTTCGGACTCGCGAACGCTCCAGCAGTATTCCAAAGGAAGATGGATCAGTGTTTCGCAGGAACCTTTGAGTTCATAGCCGTATACATAGACGACATCCTAGTGTTCAGTAAGACACTCAAGGAACATGAGAAGCATCTCAGTATAATGCTGGGAATATGTCGTGACAACGGCCTAGTACTGTCGCCAAGCAAGATGAAACTGGCAGCAACAGAAATTGACTTTCTGGGAGCAACCATTGGCGATGGGAAGATTAAGCTCCAACCTCATATCATCAAGAAGATAGCTGAGGTGGACGACGAGTCTCTTAAGACTCTGAAAGGACTAAGAAGCTGGTTGGGAGTGCTGAACTATGCCAGGAATTACATCCCAAAATGTGGAACCCTCCTCGGCCCACTCTACAGTAAGACGAGTGAGCACGGTGATAGAAGATGGCACGCTCAGGATTGGGCCTTAGTCAAAAGGATTAAAAGCCTGGTCCAGAACCTTCCGGATCTGAAGCTACCAACTGAAGAGGCCTATATAATCATTGAGACCGATGGTTGTATGGAAGGATGGGGCGGAGTCTGCAAATGGAAGCCCAACAAGGCCGACTCAGCAGGAAAAGAAGAGATTTGCGCGTACGCAAGTGGGAAATTCCCAACGGTCAAATCAACAATAGACGCGGAAATATTTGCGGTCATGGAATCCTTGGAAAAATTCAAGATATTTTACATGAACAAGGAGGAAATCACCATCAGGACCGACTGCCACGCCATCATAACTTTTTACGAAAAGTTGAACGCGAAGAAGCCTTCCCGCGTAAGGTGGCTTGCTTTCTGCGACTATATAACCAACTCCGGGGTCAGAATGAAGTTCCAACATATCAAAGGCAAGGATAATCAGTTAGCTGATAATCTCAGTCGCCTAACCCAGCTGATCACCGCAGTGAGATGGCTACCAGAGGAAATGGCAGGAATCGCGGCAGAGCTCACCAAAGAGAGGGGAATGAGCTCCGCTCTGAGCACAGTTCAGGAGAGCCTCTCAGGCTTTCTCAAAGCTGCCCTCCTCCAAGTCGAGAAGTCCTCGACTACACACCTGTCCGAGGAGCGCCCTGCTCTTTGGCAGAGATGGAAGAGTCTAGAAGACTGGCCAAACGGCTCAGAGACCGAGTCTTCAACGAAGTCAGCCGGAGCATCAGCGACACTGTATTCATCACCGGCAGGGACCTCGCAACGGCCAAAGCATACGCCACCAGGGACAACTGGTATGGCGACCTCGTCCCACTCTTGGAGAAGAGAGCAGCAGCTGCATGGAAGCTCCTCGCTGCACACGCAGAATTCTCCACATGGAAGGACGTAGACGTCTGAGAGGTGTGACGAAAAAGATGACCTCACAATTGCCAAGATC

### >endoRYNV-CU3

ACAACGGCTGTATGCTAGTCCTACCGCAGGACTTCACCTTAGTCCCTGACGTGATCAACAGATGGGAGTCCATTACAGTCAACCTCATCAACAAGATGATGTTTGACTCCCTGCAGGACAAGGCGGATTATGTGGAAAACCTCCTTGGAGAAAGGGAGAAAGAAACATGGATGACATGGAGAATGCAGTACGAAGAAGAATACAAGCAACTCCTCACCATGAGCGGGGACGTGCGAAACCTAACGGCCGCCGTCAAAAGGGTCTTCGGAGTACACGACCCACACACAGGATCTGTACACATCCAAAATCAGGCGTATGCGGAGCTAGAACGCCTCTACTGCAAAAGGACGGATGATGTGATCCCCTTCCTCTACGACTATTATCAGCTAGCAGCTAAGTCCGGAAGGATGTGGCTCGGACCCGAGCTATCTGAGAAGCTGTTCAGGAAGCTTCCACCGGAGATAGGTCCAACCATAGAGCAGGCCTACAAAGATAGATACCCAGGACTCACAATCGGAGTCTTAGCAAGGGCTAACTTCATCTTGGAGTATCTGCAGAATGTGTGCAAACAAGCTGCACTACAGAGGTCCCTCAAGAGCCTGAGCTTCTGCAGAAACATGCCAGTACCAGGATACTACGAGAAGAAGCAGTACGGCATCAGAAAGGCCAAAACCTACAAAGGTAAGCCTCACCCAACCCACGTGAAAGTCATCAAGAACAAGTACAAGCACACCTCAGGGAAGAAGTGCAAGTGCTACTTGTGTGGGATAGAAGGCCATTACGCAAGAGAATGCCCAAAGAAGGTGGTAAAACCACAGCGAGCGGCATACTTCAATGGCATGGGATTGGACGACAATTGGGACGTCGTATCAGTCGAACCCGGAGAATCAGACGATGACGAGATCTGTAGTATCTCCGAGGGAGAAAATGCTGGAGGAATGCACGAGCTGATGGCATTCAAGACTCAACTCCCCTACCCAGTGGAGTATGAAGCCAGCACACCACACCTTCTTATGCCTTGGACTCAGGTGACAATAGAAAAGAGTGAGAAACCATCCTGGAGGCGAAGGAAGGAGATCCCAAAGGCACAGCATGACTGTACTCACACCTGGAGTGATACACAGGAAGTGCCAATTGAAGGAAGGATATGCAGTATATGCAGTGACGAGACACCCCACGGGAGGAGAGTCACCTGCACCACCTGCAGCCTCAACCTCTGTCCTCTTTGCGCATATATGGATCACGGGATCAAGCTGATAGCTGCAAAGGACACGAAAGATGCAGCTAAATGGCAGTATCATAACAAAGATGAGCTCATACGACACCTCTATGAGCACAACGCTTTTCTCACAAGAAAAGTAGAGGAGTTAACCAGCCAGCTGCAGGAGTTCCACAACCGCAGGCAGGATGATCTGATCAGCCTGGCAGACGACCTGGAGGACGTGTCTATCCTAGACAACGCCTCAAAAAGAGGGAAGGAGAAGGAATCATTCCGATTTGGAACAACGATTCCCACCGATCACATCCAAAACCTTGAAAATGTAGCAAGGATGATCGAACAATGGAAGGAAACCCCCAGGGTAATCATCAAAGAAACAGCTGAGAGCAGCAGCAACACCATTGGAGCCCTCTTAGCAGAAGAAGGAATAGAGGAGCTAGCCGCAGCTGTAGACACGGCATACACTGAAATGCCAAAGGGAGGACTAAACAAGCTCTACAACACCATTGTTGAGTTTGTGATACCTCAGGAGAAAGGGGCACCCACAAGATTTAGGGTAAGGGCTGTAATCGACACAGGATGCACATGCATTAACAGCAAGAAAGTCCCCAAGGAAGCACTGGAGGAGGCGAAGTACCAGATGAACTTCGCAGGAGTAAACTCCACGGGAGAAACAAAACTCAAGATGAAGAACGGTAAGATGATCGTATCTGGAAGCGACTTCTATACACCGTACATTGCAGCTTTCCCGATGGAACTGCCAGACGTAGACATGCTCATAGGATGCAACTTCCTAAGGGCCATGAAAGGAGGAGTCAGACTTGAAGGGACAGAAGTGACAATCTACAAAAAAGTCACCACCATCCAGACTACCTTGGAGCCACAGAAAATATCTCTCCTACGCGCAGAAGCAGAAGT AGAGGAGGAGATTGAGCGCATGTACTACGCAACAGACTACTCTGAAGAGGGAGCTAGTCGTCTGAGAAATCACAAACTGCTGCAGGAGCTGAAGGAGCAAGGCTACATAGGCGAGGAGCCAATGAAGCACTGGGCAAAAAATGGAATCAAGTGCAGACTTGACATCAAGAACCCAGACATAGTAATCAGCAGCAAACCCCCGAATGCAGTCTCAAAGGAGACAAAAGCACAGTACCAGCGGCACATAGACGCCTTGCTAAAGATCGGAGTGATCCAACCAAGCAAGAGCAAGCACAGAACCGCTGCCTTCATCACAAACTCAGGGACGTCTATCGACCCTGTCACTAAGAAGGAAATCAGGGGAAAAGAAAGGATGGTGTTCGACTACAGGAGTCTGAACGACAACACCCACAAGGACCAATACACCTTGCCAGGGATCAACACCATCATATCTGCGATCGGGAATGCGAAGATCTTCAGCAAGTTTGATCTGAAGTCTGGATTCCACCAAGTGCTGATGGACGAAGAATCCATCCCGTGGACCGCATTCGTGACACCAGTAGGGTTCTACGAGTGGAAGGTAATGCCCTTCGGACTCGCAAACGCTCCGTCCGTCTTTCAAAGGAAGATGGACCAGTGTTTTGCAGGAACCTCAGAATTCATAGCCGTCTACATCGACGACATCCTGGTCTTCAGCAAGACCTTAAAGGAACATGAAAAACATCTCAGCATCATGCTTGGGATATGTCGAGACAACGGCCTGGTATTGTCACCAACTAAAATGAAGATAGCTGCAACCGAGATAGATTTCTTGGGAGCCACTATAGGTGACGGAAGAATCAAACTCCAGCCTCACATAATCAAGAAAATAGCAGAGGTGGACGATGAGTCACTCAAGACCCTCAAGGGATTGAGGAGTTGGTTGGGAGTCCTCAACTACGCGAGGAATTACATTCCAAAGTGCGGAACTCTTCTCGGCCCACTATACAGCAAGACTAGTGAGCACGGAGACAGGAGATGGCACGCATCAGATTGGGCCTTGGTCAAGAAGATCAAAAGCCTGGTCCAGAATCTCCCAGACCTAAAACTGCCCAGCGAAGAAGCCTACATGATCATAGAGACTGATGGATGCATGGAAGGCTGGGGCGGTGTTTGCAAATGGAAGCCCAACAAAGCAGACTCAGCAGGCAAAGAGGAAATCTGTGCCTACGCAAGTGGGAAGTTCCCCACGGTGAAATCAACCATTGACGCAGAAATCTTCGCGGTAATGGAGTCCTTAGAAAAATTTAAAATTTTCTACATGAACAAGGACGAAGTCACCATCCGAACTGACTGCCACGCCATCATTACCTTCTACGAGAAGTTAAACGCCAAGAAACCTTCTAGGGTAAGGTGGTTAGCTTTTTGCGACTATATAACGAACTCAGGGGTGAAGATGAAGTTCGAACACATCAAGGGTAAGGATAATCAGCTCGCTGACAACCTCAGTCGCCTTACTCAACTAATCACAGTGGTCAGATGGCTTCCGGAAGAGCTGAAGGATCTCGCGGCCGAACTAACCAAAGGAGAAGGCAAGACGCCAACGAAGAAGGAGACACAGGAAGAAATCTCCTGTTTTCTCAAAGCTGCCCTCCTCCGAGCAGAGAAATCCGCGACTACTCACCCATCAGAGCCGCACCATGCACTATGGCAGAGATGGATGAATCCCGAAGACTGGCTCTGCAACGAAGAAACAAGGTCTTCGACAGCATCGCACAAGGCATCAGCGACGCAGTGTACGTCACAGGTGTCGACCTTGCCGCCGCAAAGGCACGAGCCACCAGGGATAACTGGTACAACGACGTCACCCCGGCGCTGGAACAAAGAGCAGCTGCAGCATGGAGACTCATGGCAGCCTACTCAGACTTCGCCACGTGGAAAGACGTGAACGTCTAGTGAAGTGACGCAAGGGATGACTTCACAATTGCCAATGTCGTCATTGCTTACGACTTGGAACTTATCAGTTAGTGTCGGCAGCATCTTCCAGCTGTCAGATATTTGTGTGTAAGTGCGCCATTAGTGCGCTGTGTCAGGATAAGGAATCTTATCTCCTTATCTCTTCTTTAGCTTTAGCAAAGCTGTAAAGCAGAACTTATTAGCTGTCGATGGGGCCCAGAAAGCGCACCCGAGCTAGTAGTTGTCTGTCTTTTTGCTTAGCCTACTTCCTATATAAGGGAGTGTAGTCTGAAGGCTTAGGCATCGAGCAATCTCAATAGCCAACCTTCTCCTGTATTGTATTAACGTTCAAGAGAAATAAAGATTGTTCTTCAGTTTTCCGCAATATCTCTCAGTTTTTATGAGTTCTTAAGTGTTCAAGAGCGCATTTAAGAATTTTTGATATTGCGGAAAACTGAAGAACAATCTTTATTTCTCT

### >endoRYNV-BS

TGGTATCAGAGCTTTAGCTCTCTTCATTATGTCAGCTTAAACACCTTTTTTCGTGTCGAGAAACCCAAGTTTCGGATCTGAACCTTGAGTTCCTCTCTTTTTCAGGAGGGTAGTGAGTAAGCCCAGCTTTGCAACTTTCAAACCCCCAGGAAAACTTTCCTCACGGTATTATAAGTTTTCTGACCCTCACTAGTTCAACCCTACTGCTTGAACTGCAGACTTAGGCGTCGAAGCGAAGTACCCTTGTAGCCGTTAGCCAGGTGGGCGTTAGGCGTTGATTGGGGAAAATGGACGTAAAGGAGCAGAGGCAACTAGGCGAGAAAACTGAGGACCAGATCACCGGTCGGAAAGCCAGTAAGCGGCTAGATTGGGGCATAGCTGATGCAACCTCACGAGATCCTTTCCTTCGAAGAAGAGAGCAACTCTTGGGAACGATCTGAACGGGCGAATCGACACGACTTCCTATTCAGAAATCTCAGATCTTATCCAAGATACGAGGCTAACCAGAAATCACCTTCCTGTGATTTCCCTTGTTACCATTTCAACACCACCACCGGACCACCAGTCCACCGCACAATCTGCAAACAAAAGAACAGTGAAGACTTACCTTACCTGGTAAACACTCTGTTCGATTTATCTATAACAGGTATTCACAACCAGGCTGTCCTCGACGACAAGGTCTCGAGACTCACACAGTACCTGGTAAAGAAAGTCGGCACACTCCCGACTATCCCGGAGGATTCACCCCTCCTGGACCAGAACTCCCTGAGCTTAGATCTGCAGGCCCTAAGGGCAGATCTCAAGGAGGTCAAAGCAACTCAGTCCGCACTAAAACTCGGTTTTGAACAACTCCGAGAAGCAGTTCAATTGATCATCTCCAGGGAGAACGATCCAAAGCCAATCGAAGCCACGACTGCACTTGTGGCAGAACAGCTGAGGAAACAGTTGATTGAAGTCAGATCTGTGCTTGAGGAAACCAAGAAGGTCGTCAGATCGCTGTCCCCTAACGGATGAACCCTAAGTGGCAGGACACCGCCACAAAGGAAACCTATCAGAAGGCACTTCAAGCAACAGCATCCCTCACCTCAAACAACACAGGCCTAGGGTTCATTGAACCACACACCTTCACGGGAGGACAGTTGTCCACTAACCTAGCCAAACAGAACAACACGCTCATCCAGCTGTTGGTTCAGGCGCTAGAGAAGGTCCTCGACCTTGAACAAGCCGTCGCAAACCTGACAGCTCAGGTCACAAGGCTTGAAAAGACAGTCGCAGAGAAGGACACGGTAAAGCTACCGGAAAGCGTCCTCAACGACCTCACCAAGGAATTTGGGAAGGTCAACATCGGAAAAGGAAAAGGGATCGAAGGCACAAACACTTCCAAAGACAAGAACTTCTACGTCTGGAAGAATCCCTTCACCCAGTACAATGAGCAGAAGCCAAACAAGGCTCCAGGCTCCGCCCGCAACTGAAAGGGCCACATCTAGCTCGGACTCAGGTACCCCTACCTTGGAGGACCAGATCCGCGGCTACAGACGCTCGGCAAGGTTACGACACCAAGCCCAGCGAGCAGTTCGGAGAACCTTCAGCAGGGATTTCCGAAACACAATAGAACGGCAACTTGACCCAGATGCCGAACTCTCCCTCAGCAGAAGGAGG AGAGCAAACCTAGTACCAGCAGAGGTACTATACGCACACAACGGTTCAGAACCAGTAAACAGGGTGTATGAACACTACAGTGAGATCGGAGCTCACATAGTAGACAGGCAGCAGGACTTCAGGTACATTGAAGAATCCTCTTACCAAAGGCTAGCACGAGAAGGCATGCAGTTTATACATGTAGGCATGGCCATGGTAAGAATACAGATGCTGCACAGGACTGACGCAGGAATTTCTGCACTAGTAGTCTTTCGGGACACTAGATGGAGTGATGACAGGCAAGTCATTGGAAGCATGTCGGTAGACATGACAAGAGGTGCACAGCTGGTATATATCATACCAAATGCGATGATGTCAGTACACGACTTCTATAACCGCATACAAGTCAGCGTGCAAACCAGAGGATACGGTACAGGCTGGGCGGGAGGAGACAGCAACATGATCATCACCAGATCACTGGTGGGAAGGCTGACCAACACCAGCATGACAAATTTTGAGTACCGAATAGACCAGGTCACAGACTACCTTGCAAGCAATGGAGTCGCATGCATCCCAGGACAGAAATGGGACGTGGCCAACAGATCAGGAGAGTGGGAACTTCAGCCAAGCAGGATAGTGGCTCCACTCATGACACCTACAGAGGCAAGGCTGACCCAGAATAGAAGCGGCAGCATCAGCCTTAGGTTCACAGACTTCCGTGATCAGAGGATCGTGGAAGAAAGGCCGGCAGAAGAAGAAGGCAGGCCAGAGGAAGAAAGCACACACTATGTGCTCATGATCAGCCACAACTCAAGCTCCTATAGGACAAACCTCAAGGGAGAAAGAAGGCTGAGACAGGAAGAACTCGCTGAGTTCATGCCAGAATCAGATCTGGTAAGTCAATGGCTGAGTCAGTTATCAGCTTCAGCACACAACAGTGGAGCCTCAGACTCAGAGGACGAACCACCAAAGTTCGATGAGACTGACGAAGAAGCAGATGATGACACGTATAATCAGCGAACCTGGCAGAAAGAGGACCAGGAGAGAAGACAACTGGAACTGCAGGGGTGGAAGAAAACCAGCAGGCCAGGAATCTATGAAATGATCCCTCAGGAAGAAGAAGAAGTCTACCTCCGCTATGAGGCAGAATCAGAGGAAGAGGATCAACCCACACAGATCTTCGGAGCTACAAAGGTGGATGAACCCGAAATGGAGTACCCCACAAGGCTCGAAGAAATGATGAACAAGCTCAAAGGAGTGAGCATGGATAAGCTTTTCCCAGTGACCATGGAATCTGAGTCCAGCATAACTGGAGGAGGCTTCATACCACCAAGCCCAGTGCCCGGAGCACAAGGATACCCACCTGCAACAGGGCCAGCCTTTGGATCAACTATAGGCCCGGCAGACTTACAGGGATGGGGAGGAAGACTGCCCAGGAGCAGGTCTCCACTAGGATACGGCAGGCCGCAACAGCCATGGTCCTTGCCATCAGCCCAATCAGAAAACGGGTGTATGCTAGTGCTTCCACAAGACTTCACACTTGTTCCCGACGTGATAAACCGATGGGAATCAGTAACCGTCAACCTGATAAACAAAATGATGTTCGACTCATTACAGGATAAGGCGGACTACGTGGAGAATCTCCTTGGAGAAAGGGAGAAAGAAACATGGATGACCTGGAGAATGCAGTACGAAGAAGAATACAAGCAACTCCTCACCATGAGCGGCGATGTGAGAAACCTCACGGCTGCAGTTAAGAGGGTCTTTGGAGTACATGACCCCCACACTGGATCTGTGCATATCCAGAATCAAGCTTATGCAGAGCTAGAACGGCTGTACTGCAAAAGGACGGATGATGTGATACCTTTCCTGTACGACTACTACCAATTGGCAGCAAAGTCGGGAAGGATGTGGCTTGGCCCAGAACTGTCTGAGAAGCTATTCAGGAAATTACCTCCTGAAATCGGCCCTACAATTGAACAAGCCTACAAAGATAGGTATCCAGGCCTTACCATTGGAGTCTTGGCACGCGCCAACTTCATCCTTGAATACTTGCAGAACGTCTGCAAACAAGCAGCACTGCAGAGGTCCCTGAAGAGCCTCAGTTTCTGCAGAAACATGCCTGTGCCAGGATACTACGAAAAGGAGCAGTATGGCATTAGGAAGGCCAAGACGTACAAAGGGAAGCCCCATTCAACACACGTCAAGGTCATCAAAAACAAGTATAAGCACTCAGCAGGAAAGAAGTGCAAGTGCTATCTCTGTGGCATTGAAGGCCACTACGCTAGGGAGTGCCCAAAACAGGTGGTGAAACCACAAAGAGCCGCCTTTTTCAACGGCATGGGACTCGATGATAACTGGGATGTTGTATCTGTAGACCCAGGAGAGTCAGATGATGACGAGATCTGCAGCATATCCGAAGGAGAAGGAGCAGGAGGAATGAACGAGCTCCTAGCCTTTAAAACACAACTGCCATACCATGTGGACTACGAGGCCACAACATCATCACAGCAGTTAATGCCATGGATACAGGTCACTGTATCAAGAAGTGAAAAACCATCCTGGAGACGACGGAAGGAAGTCCGAAAGGAACAGCAGGATTGTTCTCACCAGTGGAGTGACACGAAGGAAGTGCCACTAGAGGCGAGAGTCTGCAGCATTTGCAGTGATGAAACTCCATTAGGCAGGAGGATCACCTGCGAAACCTGCAATCTGAATCTGTGTCCCATCTGTGCATATATGGATTACGGAATCCAATTAGTAGCAGCTAAAGACACAAGGGACGCTGCAAAATGGCAGTACAACAACAAAGACGAGCTCATCCGCCAACTATATGACCATAACGCTTTTCTAAGCAGGAAAGTCGATGAACTCACCAGCCAGCTGCAACAATACAGAGATCGCAAACCTGAAGATCTCATCAGCCTCGCAGATGACCCAGAGGACATGTCCATACTGGACGATGCCTCAAAAAGACGGGGGAAGGAGCATGAAATCTTCCAGTTCGGAACGACATTGCCCACGGATCACCTCCAGAACATCGAGAATGTCGGGCGAATGATCGAACAGTGGAAAAACAACACCCCCAGAGTCACCATCAAAGAAATAATAGGTGACGAAATGCCTAGCAGCAGCAGCACAATAGGAGCACTCCTCGCAGAAGAAGGAATCGAGGAACTCGCTGTAGCAGTAGACACGGCGTATACTGAAATGCCCAAGGGAGGACTCAACAAGCTCTACAACACGGTTGTGGAATTTGTCATCCCCCAAGACAAGGGAGCGCCAACAAGATTTCGGGTAAGAGCAGTCATTGATACAGGATGCACCTGCACCTGCATAAATAGCAACAAAGTCCCGAAAGAAGCCATGGAAGAAGCAAAGTACCAGATGAACTTTGCAGGAGTAAATTCAGCGGGTAGTACAAGGATGAAGATGAAGAGCGGAAAAATGATTGTATCTGGGAGTGACTTCTACACCCCGTACATAGCCGCCTTTCCCATGAACCTGCCTGAAGTAGACATGCTCATAGGCTGCAACTTCCTACGAGCCATGAAAGGAGGGGTCAGATTGGAAGGAACAGAGGTGACAATCTACAAGAAAGTCACGACTATTCAGACGACCCTCGAACCACAGAAAATCTCCCTGATGAAAGCAGAGATGGAAGCTGAAGAGGAGATTGAAAGGATATACTACGCCAGTGATCATTCGGAAGAAGGAATGAACAGGCTTAGAAACCACAAGCTGCTCCAGGAGCTTAAGGAGCAAGGATACATAGGGGAAGAGCCCATGAAGCATTGGGCCAAGAATGGAATCAAGTGCAAACTTGATATCAAAAACCCCGACATCGTCATCAGCAGCAAGCCGCCTGATGCTGTATCCAAAGAAACAAAGGCCCAGTACCAGCGACACATTGATGCTCTGTTAAAGATAGGGGTCATCCGGCCCAGCAAAAGCAAGCATCGCACCGCGGCCTTTATCACACATTCGGGCACAAGCATTGACCCGATAACCAAGAAGGAGATACGGGGAAAGGAAAGAATGGTCTTTGACTACCGAAGCCTGAACGACAACACCCACAAGGATCAGTACACCTTGCCAGGCATAAACACAATCATCTCCGCCATTGGAAACGCGAAGGTCTTCAGCAAGTTCGACCTGAAATCAGGATTCCACCAAGTCCTCATGGACGAGGAATCAGTACCATGGACCGCGTTTGTCACACCCGTAGGGTTCTACGAGTGGCTCGTCATGCCATTCGGCCTCGCAAATGCACCTGCAGTATTCCAAAGAAAGATGGACCAGTGCTTTGCAGGTACTTCAGATTTTATTGCTGTGTACATTGACGACATACTCGTTTTCAGCAAGACAATCAAAGAACATGAGC GTCACCTGAGCATCATGTTGAGCATCTGTCGAGACAACGGCCTGATCCTCTCGCCTAGCAAAATGAAGATCGCCGCCACGGAAATTGATTTCCTCGGGGCAACCATAGGTGACGGAAAGATCAAGCTGCAACCGCATATCATTAAGAAGATTGCGGAAGTCGATGATGAATCCTTAAAGACCCTCAAAGGATTAAGGAGCTGGCTAGGAGTCCTCAACTACGCCAGAAACTACATTCCCAAATGCGGAACCTTGCTTGGGCCTCTGTATAGCAAGACAAGTGAACATGGGGATAGAAGGTGGCACGCACAGGATTGGGCTTTGGTTAGAAAAATAAAGGCCTTGGTCCAAAATCTGCCAAACCTGCAATTACCCACTGAGGAAGCCTACATCATAATTGAGACTGATGGATGTATGGAAGGCTGGGGAGGCGTATGCAAGTGGAAGCCCAACAAGGCTGATTCACCAAGCAAAGAAGAAATATGCGCCTATGCAAGCGGTAAATTCCCTACCGTAAAATCTACCATTGATGCGGAAATCTTTGCTGTCATGGAATCATTGGAGAAATTCATAATCTTCTACATGAACAAAGATGAGATCATCATCCGCACAGACTGCCACGCCATCATCACGTTCTACGAAAAGTTGAACGCAAAGAAACCTTCTCGGGTAAGGTGGCTAGCCTTTTGTGACTATATAACTAACTCTGGTGTCAAGATGATATTCCAACATATCAAAGGGAAGGACAACCAGCTCGCGGATAATCTAAGCCGCCTCACACAGCTCATCACTGCAGTCAGATGGCTCCCAACAGAAATGGCAGGAATCGCCAGCGAACTCACCAAGGAGAGAGCTCCGAGTCCAGCAATGGACGAGGTACAGAAGAACCTGTCCGGCTTTCTAGAAGCTGCCCTCCACCAAGTCGAGAAATCCTCGACTATGAACCATTCCGCAGAGCACCATGCTCTATGGCGGACATGGAGGAATCTCGAAGAGCTGCCAAACAGCTCCGGGACAGAGTTTTCAACGAGGCAGGAAGACAGATCAGCGACATCGTCTTCATCACAGGAAAGGACCTCGCAGCAGCCAAGGCATACGCTACCAGGGACAATTGGTACGGCGACCTCGTCGGCCTCCTGGAGAAACGCGCCGAAGCGGCGTGGAAGCTGTTAGTCGCATACTCCGAGTTCTCTACGTGGAAAGTGGATGTCTAGAAGACATGACGTAGGAGCTCACCTCAGTAATTGCCGGGACGTCACTGCGTGCGTCACGGAACTTATCTTTTAGTGTCGGCAGCACGTATTAGCTGTCACGCATTTTGTAAGTGCGCCATTAGTGCGCGTGAGTCAGGATAAGGAATCTTATCCTTTGCTTTAGTTAGTGGCAAGCTGTAAAGCAAGCATTATTAGCTGTCGATGGGGCCCCAAGCGCACCCGGGCTAATTTTCTCTTGTCTTTTAGCATAAGCCCCCTTCCTATATAAGGAAGCTAAGTTA GAAGGCTTAGGCATCGAGCAACCTCAATAGCCAACCTTCTCTTGTAATATCAGTATTCAAGAAATTAGTATTCAAGAAATTAGTTG

### >endoRYNV-PH1

TGGTATCAGAGCTTTAGCTCTCACCATGGTAGCTTAAACACCTTTTTCTTGTCGAGAAACCCAAGTTTCAGATCTGAACCTTGAGCTTTCTCTCTTTTTCTAGGGCGAAAGGAGTGTGACCCTGTTCAAAACCTTACCTGATCAAACCCCCATGAAAACTTTCCTTACGGTACTATAAGTTTTCTGTCTATCACTAGTCCAAACCTACTGCTCAAACTGCAGGTCTAGGCGTCGAAGCGAAGTACCCTTGTAGCCGTTAGCAGGAGGCGTTAGGCATTGATTGGGGAAAACTGACGTAAGAAAGCAGCAGCAACTAGGCAAGGAACTTGACAGACAGATCACCGGCCGGAAAGCTAGTAAGCGGCTAGATCTTGGTGGTTTTGATGCAACCACACGAAATCTCTGCCTTCGAAGAAGAGAGTAGCTCCTGGGAAAGATCTGAACGGGCGACTCGGCAAGACTCTTTATTCAAGAATCTCAAAAACCTATCCACGTTGGGAGGCGAATCAAAAGACACCCTCTCTCGATTTCCCGTGCTACCACTTCAACACAACATCTGAAGAAACAGCTCATCGAGGTCAAGACAGTCCTCGAAGAGACCAAGAAGATCGCTAGGTCTTTATCCCCCGACGGATGAACCCTAGGTGGCAGGATACTGCAGCCAAGGAGACCTACATCAAAGCGATCCAAGCTACCTCATCGCTTACCTCCAGCAACACTGGTCAGGGTTTCATCGAACCCCACATCTACACAGGAGGACAGCTATCCACTAACCTAGCAAAACAGAACAACACTATCATCCAGTTGTTGGTTCAAGTGCTAGAGAAGAACCTCGACCTTGAGCAGGCAGTCGCCAACCTCACAGCTCAGGTCACGAGACTAGAAAAGGCCGTCGCTGACAAAGACACGGTCAAGCTCCCTGAGAGTGTCCTGAACGACCTCACAAAAGAGTTCGGGAAGGTTAACTTAGGGAAGGGCAAGGGGATAGAAGGAACCGTCTCTTCTAGAGACAAGAACTTCTACGTCTGGAAGAACCCCTTCAACCAGTACAATGAACAGAAGCCACACAAGGCTCCAGGCTCCGCCCGCAACTGAAAGGGCCACGAGCAGCTCGGACGCAGGAACCCCCACTCTGGAGGACCAGATCCGAGGCTACAGACGCTCTGCACGGTTACGACACCAGGCGCAGCGCACCATGAGAAGGACCTTCAGCAGGGACTTCCAACGGCAACTAGATCCGGATGCCGAGCTCTCTCTCAGCAGAAGGGGAAGAGCAAACCTGGTACCAGCAGAGGTACTACACACACTGAAAGAAGAAGGACCATCCGAGCCAGAAGGAAGGCCAGAAGGAGAGGACGAGAGCACACATTATGTGCTAATGTTCAACCATTCCAGCCCCAGATGGGATACGCTCGGACAGCCAAGCGGAAAATATGACTACATGGTACGGTATGATGCACCGGAACCAACCGCATGGCCGACAACCAATATAGGATGGGATGACGACCCACCAAAGCCACCAAGCCCTAAAGGATCTTTTGAGATCAACCTAAGAGGCGAAAAACGACTAAAAGAGAAGGAACTCTCAGAGTTCACTCCTGAAACTGACCTAGTCAGTCAGTGGTTGAGTCAGCTCTCCAACTCTGCACACAACAGTGGAACTTCGAGCTCTGAAGAAGAGCCACAGTTCGACGAGGCAGACGACGAGAACGACGAGTACAACCAGCAAACCTGGCAACGAGAGGACCAAGAAAAGAGAGACCTGGAACTACAAGGGTGGAAACCTACTGGTAGACCAGGAATTTACGAAATGATCCCCGAAGAAGAAGAAGAAGTCTACCTCAAATATGAGGCAGAAGACGAAGAGGAGGATCAGGAGCTTCAAGTGATTGGAGCCACCACCATAGAAGAGCCAGAAATGGAATACCCAACAAGGCTCGAGGAAGTTATGGGCAAGCTCAAGAACGTGAGCATGGAGAAGCTGTTCCCAGTCAGTGGGATGGACAGCGAATCCAGCATAACAGGTGGAGGATTTATCCCACCTAGCCCGGTGCCAGGCGCACAAGGATATCCCCCAGCAACAAGCGCATCAGCGTCCACAATAGGACCAGCAGACATGCAGGGATGGGGAGGAAGAATGCCTAGAAGCAGATCACCTCTGGGCTATGGCAGACCTCAACAACCTTGGTCATTGCCATCTGCACAGTCAGACAATGGCTGTATGCTAGTCCTGCCACAGGACTTCACACTAGTCCCGGACGTCATCAACAGATGGGAGTCTATCACAGTCAATCTCATCAATAAGATGATGTTTGACTCTCTCCAAGACAAGGCGGACTACGTCGAAAATCTCCTAGGCGAAAGAGAGAAGGAGACATGGATGACATGGAGGATGCAGTACGAGGAAGAGTACAAGCAGCTCCTAACCATGAGCGGGGACGTGAGAAATCTCACCGCCGCAGTAAAACGGGTCTT CGGAGTACATGACCCACACACAGGATCAGTGCACATCCAGAACCAAGCATATGCGGAGCTAGAGCGGCTCTACTGCAAACGAATGGATGATGTGATCCCCTTCCTCTATGACTACTACCAGCTAGCAGCCAAATCAGGAAGGATGTGGCTCGGACCCGAGCTGTCGGAGAAGCTATTTAGAAAGCTCCCCCCGGAAATAGGCCCAACAATAGAACAGGCCTATAAAGACAGATACCCAGGGTTGACGATAGGAGTCCTAGCAAGGGCCAACTTCATCCTGGAGTATTTACAAAACGTATGCAAGCAGGCAGCCTTGCAACGATCTCTCAAGAGCCTGAGCTTCTGCAGGAACATGCCAGTACCAGGGTACTATGAGAAGAAGCAATATGGCATAAGGAAGGCTAAAACCTACAAAGGTAAGCCACATCCAACCCACGTGAAAGTGATCAAGAACAAGTACAAGCACACCCAAGGGAAGAAGTGCAAGTGCTACCTATGTGGGATAGAAGGCCACTACGCCAGAGAATGCCCGAAGAAGGTGGTCAAACCTCAAAGAGCGGCATACTTCAACGGCATGGGCCTAGATGATAACTGGGATGTCGTATCTGTCGAACCAGGAGAGTCCGACGACGACGAAATCTGCAGTATCTCAGAAGGAGAGAACGCTGGTGGAATGCACGAGCTGATGGCATTCAAGACCCAACTCCCGTACCCAGTGGAGTATGAAGCCAGCACACCACAGTTCCTGATGCCTTGGACACAGGTAACAGTGGAGAGAAGTGACAAACCTTCCTGGAGAAGAAGGAAGGAAATCCCAAAGGTACAGCAGGACTGCACGCACACCTGGAGCGACACGCAGGAGGTACCTATAGAAGGCAGGATATGCAGCATATGCAGCGATGAGACACCTCACGGACGGAGAGTGACCTGCACAACATGCAGCATCAACCTGTGCCCCCTTTGCGCGTATATGGATCACGGGATCAAGCTAATAGCCGCAAAGGACACCAAAGACGCAGCCAAATGGCAATATCACAACAAGGACGAGCTGATTCGTCACCTGTACGAGCACAATGCTTTCCTCACCAGGAAGGTAGAAGAACTCACCAGCCAGCTGCAAGAATTCCACAGCCGCAGACCTGAAGACCTGATCAGTCTGGCGGACGACTTGGAGGACGTGTCCATTCTGGACAACGCCTCAAAAAGGGGGAAGGAGAAGGAATTGTTCCAATTTGGAACGACAATTCCCATTGACCACATCCAAAACCTGGAAAACGTGGCCAAAATCATCGAACAATGGAAAGACACCCCCAGGGTTGTGATCAAAGAGACACCTGAAAGCAGCAACAACACCATTGGAGCCCTCCTAGCCGAAGAAGGGATAGAAGAACTAGCTGCAGCAGTGGACACAGCCTACACGGAGATGCCAAAAGGAGGCCTCAACAAACTTTACAACACCATTGTAGAGTTTGTAATACCACAAGAAAAAGGGGCACCCACCAGATTCAGGGTACGAGCTGTAATAGACACAGGATGCACCTGCACATGCATCAACAGCAAGAAAGTCCCCAAAGAGGCACTAGAAGAGGCAAAGTACCAGATGAACTTTGCAGGAGTAAACTCAACGGGAGAAACAAAGCTCAAGATGAAAAATGGTAAGATGATCGTCTCTGGGAGCGATTTCTACACACCATACATAGCAGCTTTCCCGATGGAGCTACCAGACGTAGACATGCTCATAGGATGCAACTTCCTAAGGGCTATGAAAGGAGGCGTTCGTCTTGAAGGAACGGAGGTGACGATCTATAAAAAGGTTACCACCATCCAGACCACCTTAGAGCCTCAGAAGATATCCCTCCTCCGCGCAGAAGCTGAAGCAGAAGAGGAGATGGAGCGCGTGTACTACGCGAATGACTATTCCGAAGAAGGAATCAGTCGCCTGAAGAACCATAGGCTGCTACAGGAGCTGAAGGAACAAGGCTACATTGGCGAGGAACCAATGAAGCACTGGGCCAAAAATGGTATCAAGTGCAAGCTTGATATTAAAAACCCAGATATAGTGATCAGCAGCAAGCCCCCTGACTCAGTGTCAAAAGAGACAAAGGCCCAATACCAGCGGCACATTGACGCCCTACTCAAGATAGGAGTCATACAGCCTAGTAAGAGCAAGCACAGGACCGCGGCCTTCATCACACATTCGGGTACGTCCATTGACCCGATCACAAAGAAAGAAGTCAGAGGAAAAGAAAGGATGGTGTTCGACTACAGAAGTCTAAACGACAACACCCACAAAGACCAATACACTTTGCCTGGGATCAACACCATCATCTCAGCGATTGGCAATGCAAAGATCTTCAGTAAGTTTGATCTGAAGTCTGGATTCCACCAGGTGCTGATGGACGAAGAATCCATCCCATGGACTGCATTTGTCACGCCAGTAGGGTTCTACGAGTGGAAGGTCATGCCCTTCGGACTCGCAAACGCTCCGGCCGTCTTCCAGAGAAAGATGGACCAGTGTTTTGCAGGAACCTCAGAGTTCATCGCCGTCTACATCGACGACATCCTGGTCTTTAGCAAGACCTTGAAGGAGCACGAAAAGCACCTTAGCATCATGCTAGGGATCTGTCGAGACAACGGCCTGGTATTGTCACCCAGCAAAATGAAGCTTGCAGCAACCGAGATCGATTTCTTGGGAGCTACCATAGGTGACGGAAGGATCAAGCTCCAGCCTCACATAATCAAGAAGATTGCAGAGGTGGACGACGAGTCCCTAAAAACCCTCAAAGGGTTAAGAAATTGGTTGGGAGTTCTCAACTATGCCCGAAACTACATCCCGAAGTGTGGAACACTCCTCGGCCCACTATACAGCAAAACTAGTGAGCATGGAGACCGTAGGTGGCATGCCTCAGATTGGGCCTTAGTAAAGAAGATCAAAGGCCTGGTCCAAAATCTCCCAGACCTGAAACTGCCCACAGAAGAAGCCTACATGATCATCGAAACCGATGGATGCATGGAAGGTTGGGGAGGAGTCTGCAAGTGGAAGCCCAACAAAGCAGACTCAGCTGGAAAAGAAGAAATCTGCGCTTACGCAAGTGGGAAATTCCCCACAGTAAAATCCACGATTGACGCAGAAATCTTCGCGGTAATGGAGTCCTTGGAGAAGTTCAAAATCTTCTACATGAACAAGGACGAGGTCACAATCAGGACTGACTGTCAGGCCATCATCACCTTCTATGAGAAGCTGAACGCTAAGAAACCTTCACGGGTAAGGTGGTTAGCTTTTTGCGACTATATAACAAACTCAGGGGTGAAGATGAAGTTCGAACACATCAAAGGCAAAGACAATCAGCTTGCCGATAACCTCAGCCGATTGACCCAACTCATCACCCTGGTCAAATGGCTTCCAGAAGAACTGAAGGATCTCGCGGCCGAGCTAGCCAAGAAAGAAGGCAAGACCTCCCTGAAGGGAGAGGTGCAGGAGGAGATCTCCTGTTTTCTCAGAACTGCCCTCCGCCGAGCAGAGGAATCCGCGACTACTCGCCCATCAGAGCCGCGCCATGTACTATGGCAGAGATGGATGAATCCCGAAGACTGGCCTTACAACGAAGAAACAAAGTCTTCGATGACATCGCACAGAACATCAGCGATGCGGTCTACATCACCGGCATCGACCTCGCAGCGGCAAAAGCAAGAGCTACCAGGGACAACTGGTACAATGACGTCACACCCGCACTGGAGCAAAGAGCAGCTGCAGCATGGAGACTCATGGCAGCCTACTCAGACTTTGCCACGTGGAAGGACGTAAACGTCTAGTGAAGTGACGCAAGGGATGACTTCACAATTGCCAATGTCGTCATTGCTTACGACTTGGAACTTATCAGTTTGTGTCGGCAGCATCTATTAGCTGTCATACTTTATGTAAGTGCGCCAGTAGTGCGCTGAGTCATAAGGATAAGGAATCTTAGCTCCTTATCGTCCTTTAGCTTTAGCAAAGCTGTAAAGATGAATTTATTAGCTGTCGATGGGGCCCAGAAAGCGCACCTGAGCTGATATTTTCTATCTTTTTGCTTAGCCTACTTCCTATATAAGGGAGTATAGTCTGAAGGCTTAGGCACAGAGCAATCTCTTTAGCCAACCTTCTCTTGAGTTGTATTAA

### >endoRYNV-PH2

TGATACCAGAAACAAAGAATTACCATGGTCGCTTAAATTTTCTTTTCTTGTCGAGAAACCCAAGTTTCAGACCAGAACCTTGAGTTGTCTCTCCTTTTTTACAGGGGAGGAGGGAGTACTGAGTGTTGCTAAAATTCTGCAAATTTCAAACCCCCATGAAAACTTTCCTCACGGTATTATAAGTTTTCTGACCCTCACTAGTTCAAACCTACTGCTGAAACTGTAGACTTAGGCGTCGAAGCGAAGTACCCTTGTAGCCGTTAGCAGAGTGGCGTTAGGCGTTGATTGGGGAAAACTGACGTAAAGGAACAGCTGCAACTAGGCAAGAAAACTGAAGGTCAGATCACCGGCCGGAGAGCTAGTACGCGGCTAGATCTGGGCAGATCTCTTGATGCAACCTCACGAAATCTCATCTTTTGAAGAAGAGAGCAACTCTTGGGAAAGGTCTGAACGGGCGTATCGACAAGACTTCTTATTCAGGAATCTAAGATCCTACCCTCGTTACGAAGCAAACCAGAAGTCACCTTCCTGTGATTTCCCTTGCTATCACTTCAACACAACAACCGGACCACCAGTCCACCGCACTCTCTGCAGACAGAAGAACAGCGAAGATTTACCCTTCCTGGTAAATACTCTGTTCGATCTTAGTATCACCAAGATTCACAACCAAGCGGTCTTAGACGATAAGATCTCAAGACTCACCCAGTACTTGGTGGCAAAAGTCGGCACTCTACCGACAATCCCGGAGGAATCACCCCTCCTGGACCAGACAGCCATCTCTCTAGATCTTCAAGCCCTCAAGGCAGATCTAAAAGAGATCAAGGCTACCCAGTCATTCTTGAAGCTAGGCTTTGCACAGCTCCAAGAAGCGGTACAGCTGATCATCACAAGGGAGAACGATCCCAAACCGATTGAGGCAGCTACAGCTCAGGTGGCTGAACAACTGAGGAAGCAGCTTATCGAGGTTAAAGCCGTCCTAGAGGAAACAAAGAAGATCGCGAGATCTCTGTCCCCCGACGGATGAACCCCAGGTGGCAAGACACCGCCGCAAAAGAAACCTACATAAAAGCTATCCAGGCTACCTCATCTCTTACATCCAACAACACCGGGCTAGGGTTCATTGAACCTCACACCTATACCGGAGGACAACTGTCCACAAGCCTAGCAAAACAAAACAACACGCTCATACAGCTGTTGGTTCAGGTGCTAGAGAAGAATCTCGACCTTGAACAGGCCATCGCGAACCTGTCCGCACAGGTCACGAGACTTGAGAAGACCGTCGCAGAGAAAGACACGGTCAAACTCCCAGAAAGTGTCCTCAACGACCTCACTAAGGAGTTCGGAAAGGTCAACCTTGGGAAAGGAAAGGGGCTAGAAGGAGCAGTCTCATCCAGAGACAAAAACTTCTACGTTTGAAAGAATCCCTTCAATCAATACAATGAGCAGAAGCCAAACAAGGCTCCAGGCTCCGCCCGCAACTGAAAGGGCAACAAGCAGCTCGGACTCAGGCACCCCCACTCTGGAGGACCAGATCCGAGGATACAGGCGCTCTGCAAGGTTACGACACCAAGCACAGAGAGCGGTACGAAGGACCTTCAGCAGGGACTTCAGGAACACCATAGAAAGGCAGCTAGATCCGGACGCCGAGTTATCACTCAGCAGAAGGAGAAGAGCAAACCTAGTACCAGCAGAGGTACTCTACGCTCACAACGGTTCAGAACCTGTGAATCGTGTGTATGAGCACTATAGTGAGCTCGGGGCTCATATAGTAGACAGGCAACAAGACTTCCGATATATCGAGGAAGCATCCTACCAGAGACTAGTCAGAGAAGGCATGCAATTCATACATGTCGGCATGGCAATGGTTAGGATCCAAATGTTGCACAGGACAGATGCAGGTATATCTGCGTTGGTGGTGTTCAGAGACACTAGATGGAGCGATGACAGGCAGGTCATCGGCAGCATGTCCGTAGACATGACCAGGGGAGCACAATTGGTATACATAATACCAAACGCAATGATGTCAATACACGATTTTTACAATCGTATACAGGTCAGCGTGCAAACCCGAGGCTACGGGACAGGATGGGCTGGAGGAGACAGCAATATGATCATCACTAGATCACTGGTGGGGCGTCTCACCAACACCAGCATGACAAACTTCGAGTACCGGATAGATCAAGTCACAGACTATCTAGCAAGCAACGGAGTCGCTTGTATACCCGGACAGAAGTGGGATGTGGCCAACAGGTCCGGCGAATGGGAGCTACAGCCCAGCAGGATTGTTGCACCACTCATGACACCAACAGAGGCGAGACTCAGCCAGAACAGGAGTGGCAGTATAAGTCTCAGATTCACCGACTTCCGAGATCAGAGGATCGTGGAAGAAAGGCCAGTCGAGGATGAGGGCAGGCCAGAAGGTGAAGAGGAGAGCACGCACTACGTGCTCGTCTTCAGACATAACTCCTATGAGACCAACCTCAGAGGAGAAAGGAGGCCAAGGCAGAACGAACTTTCAGAGTTCACACCAGAGACAGATTTGGTGAGCCAATGGCTAAGCCAGCTATCCGCATCAGCACACAACAGTGGAGTGTCAAGCTCAGAAGAAGAACCGCCTAGGTTCGATGAAACTGATGAAGACAGTGATGGCACATACAATGAGAAAACCTGGCAGAAGGAAGACCAGGAAAGGAGAAATCTGGAGCTGCAAGGATGGAAGAAGACCTGCAGACCAGGCATATACGAACTGATCCCAGAGGAAGAAGAGGAAATCTACCTCAGATACGAGGCAGAAGACGAGGATCAGGAGGTACAGGTACTAGGGGCAACAACCATGGAGGAACCAGAAATGGAGTACCCCACCAGACTGGAAGAGGTTATGGGCAAGCTAAAGAATGTCAGCATGGAGAAACTTTTTCCAGTAAGTGGCATGGATAGCGAGTCCAGCATCACAGGTGGAGGATTCATACCACCAAGCCCAGTACCAGGAGCGCAGGGATACCCACCTGCCACAGGAGCGTCTGCTTCAACCATAGGGCCAGCAGATCTGCAAGGATGGGGAGGACGATTGCCAAGGAGCAGATCGCCGATAGGATATGGCCGCCCACAGCAGCCATGGTCCTTACCATCAGCCCAGTCTGACAACGGCTGTATGCTAGTCCTACCTCAGGATTTCACCTTAGTTCCCGATGTGATCAACCGATGGGAGTCTATTACTGTCAACCTCATCAACAAAATGATGTTTGATTCCCTACAGGATAAGGCGGACTATGTAGAAAATCTCTTGGGAGAAAGAGAAAAGGAGACATGGATGACTTGGAGGATGCAGTACGAAGAAGAATATAAGCAACTCCTCACCATGAGCGGAGACGTGAGGAATCTCACTGCCGCAGTCAAAAGGGTCTTTGGCGTACATGACCCTCATACTGGATCAGTCCACATACAGAACCAGGCGTATGCAGAACTGGAGAGACTGTACTGTAAGCGGACAGACGACGTGATCCCCTTCCTCTACGACTACTACCAGCTAGCAGCTAAGTCTGGAAGGATGTGGCTCGGACCAGAGCTATCAGAAAAACTTTTCAGAAAGCTTCCACCTGAGATAGGGCCAACTATTGAACAGGCCTACAAAGACAGGTACCCAGGTCTCACCATTGGAGTCTTGGCAAGGGCCAACTTCATCCTGGAATACCTACAAAACGTCTGCAAGCAAGCAGCGTTACAGAGGTCCCTGAAGAGCCTCAGTTTTTGCAGAAACATGCCGGTGCCTGGATACTATGAGAAGAAACAGTATGGCATCAGGAAGGCAAAAACTTACAAGGGAAAGCCTCACCCAACGCACGTAAAGGTGATCAAAAACAAGTACAAGCACACCTCCGGGAAGAAGTGCAAGTGCTACCTTTGCGGAATAGAAGGCCATTACGCCAGGGAATGTCCAAAGAAGGTGGTAAAACCACAAAGGGCGGCGTATTTCAATGGCATGGGCTTAGATGACAACTGGGATGTTGTATCCGTAGAGCCAGGAGAGTCAGATGACGACGAAATCTGCAGCATCTCCGAAGGAGAGAACGCTGGGGGAATGCATGAACTCATGGCATTCAAAACTCAACTCCCGTACCCAGTCGAGTACGAAGCCAGCACATCACAACAGTTCCTGCCATGGATACAGGTAACTGTCGAGAAAAGTGATAAGCCCTCTTGGAGGAGAAGA AGAGATATCCCACAGGCACAAAAGGACTGCGCTCACACTTGGAGCGACACACAGGAAGTGCCGATAGAAGGGAGGATATGCAGCATTTGCAGTGATGAAACGCCTCATGGAAGGAGAGTCACCTGCACGACCTGCAGCCTAAACCTGTGTCCAATTTGCGCATATATGGATCATGGAATTAAGCTGATCGCCGCAAGGGACACCAGAGACGCAGCAAAGTGGCAGTACCACAACAAAGATGAACTCATCAGGCACCTCTATGAACACAATGCCTTCCTAACCAGGAAAGTGGAGGAACTGACGACCCAGCTGCAAGAATTCCAAAACCGCAAGCCTGAAGACCTAATCAGCTTGGCGGATGATATGGAGGACGTGTCCATTCTGGACAACGCCTCAAAAAGGGGGAAGGAGAAGGAATCTTTCCAATTCGGAACGACGATTCCCATCGATCACATCCAGAATCTTGAAAACGTAGCAAGAATGATCGAACAATGGAAGGAGACCCCCAGAGTCACTATCAAGGAAACAGCAGAAAGCAGCAACACAATAGGAGCCCTCCTAGCAGAAGAAGGGATTGAAGAGCTAGCTGCAGCTGTTGATACGGCATACACAGAAATGCCAAAAGGAGGTCTGAACAAACTATACAACACCATTGTTGAGTTTGTAATACCTCAGGAAAAAGGGGCACCCACAAGATTCAGGGTTAGAGCAGTCATAGACACAGGATGCACCTGCACATGCATCAACAGCAAGAAGGTCCCTAAGGAAGCCCTGGAAGAAGCAAAGTATCAGATGAACTTTGCAGGAGTAAATTCCACTGGAGAAACCAAGCTGAAAATGAAGAATGGCAAGATGATAGTATCAGGAAGTGATTTCTACACGCCATATATTGCAGCTTTCCCAATGGAATTACCAGATGTGGACATGCTCATTGGCTGCAACTTCTTGCGAGCCATGAAAGGAGGAGTCAGGCTGGAAGGAACGGAGGTAACCATCTACAAGAAGGTTACAACTATCCAGACGACTCTAGAACCACAGAAGATCTCTCTACTCCGTGCAGAAGCTGAAGCAGGAGAAGAAATCGAGAGACTTTACTACGCCAACGACTACTCTGAAGAAGGAGTAAACAGGCTGAGAAACCATAAGCTGCTACAAGAGCTCAAGGAGCAGGGTTACATTGGGGAAGAACCAATGAAGCACTGGGCGAAGAACGGAATCAAGTGCAAGCTTGATATCAAGAACCCAGATATAGTCATCAGCAGCAAACCTCCGGATGCAGTCTCAAAAGAGACGAAGGCACAGTACCAGAGACACATCGATGCTCTATTGAAGATCAAGGTAATCCAACCCAGCAAGAGCAAGCATCGAACTGCCACCTTCATCACAAACTCGGGTACAACCATAGACCCGATAACAAAGAAGGAGATCCGAGGAAAAGAGAGGATGGTCTTTGATTACAGAAGCCTCAACGATAACACCCACAAGGATCAGTACACCTTGCCTGGGATTAATACCATCATCTCCGCAATTGGCAACGCAAAAATCTTCAGCAAATTTGACTTGAAGTCTGGCTTCCACCAGGTACTTATGGATGAGGAGTCCATCCCTTGGACCGCGTTTGTCACACCAGTAGGTTTCTACGAATGGAAGGTAATGCCTTTCGGACTCGCGAACGCTCCTGCAGTCTTCCAGAGGAAGATGGACCAGTGCTTCGCAGGAACCTCTGAGTTCATAGCCGTCTACATCGACGACATTCTGGTCTTCAGTAAGACGCTGAAGGAACATGAGAAGCATCTCAGCATCATGCTGGGAATATGTCGAGATAACGGCCTGGTACTGTCACCCAGCAAAATGAAGCTCGCCGCAACAGAGATAGATTTCTTGGGAGCAACCATTGGTGACGGAAAAATCAAGCTCCAGCCTCACATCATCAAGAAGATTGCCGAGGTGGACGATGAATCCCTAAAGACCCTCAAAGGGCTGAGAAGCTGGTTGGGAGTTCTGAACTATGCCAGGAACTACATCCCCAAGTGTGGGACACTCCTTGGCCCACTCTACAGCAAGACGAGTGAGCACGGGGACAGAAGATGGCATGCATCGGATTGGGCCTTAGTCAAGAAAATCAAAAGCCTGGTCCAAAACCTCCCAGACCTCAAACTGCCCACTGAAGAGGCCTATATGATCATCGAGACAGATGGTTGTATGGAAGGATGGGGCGGAGTCTGTAAATGGAAGCCCAACAAGGCAGACTCAGCAGGAAAGGAAGAAATCTGCGCATATGCAAGTGGGAAATTCCCAACGGTCAAATCAACTATTGACGCAGAAATCTTCGCGGTCATGGAATCCTTGGAAAAATTCAAGATATTTTACATGAACAAGGATGAGATCACCATCAGGACCGACTGCCACGCCATCATAACCTTTTACGAGAAGTTAAACGCCAAGAAGCCTTCACGCGTAAGGTGGTTAGCTTTTTGCGACTATATAACAAACTCCGGGGTGAGGATGAAGTTCGAACATATCAAAGGCAAAGATAATCAATTAGCTGATAATCTCAGTCGCCTAACCCAACTCATCACCGCAGTAAGATGGCTTCCAGAGGAAATGGCAGGAATCGCGGCAGAACTAGTCAAAGACAGGGAAGAGTCCCCCGCGATGCAGAAGGTACAGGAGAGCCTCTCAGGCTTTCTCAAAGCTGCCCTCCTCCAAGTCGAGAAATCCTCGACTACACGCCCATCAGAGCCGCACCATGCTCTCTGGCAGAGATGGACGAGTCTAGAAGACTGGCCAAACAACGCCGAGACAAAGTCTTCGACGACGCAGGGCAAAACATCTGCGACACGGTCTACATCACCGGTGTCGACCTCGCCGCCGCCAAGGC

### >epiRYNV-BS

TGGTATCAGAGCTTTAGCTTTCACCATGGTAGCTTAAACCCCCCCCATCTTGTCGAGAAACCCAAGTTTCAGATCCGAACCTTGAGTTTGCTCTCTTTTCGAAGAGGAGAGGGAGTGTGACCTTACTCAAAACTTTTGCGGATCAAACCCCCACGAAAACTTCCTTCACGGTATCATAAGTTTTCTGTCCTTCACTAGTCCAAACCTACTGCTCAAATCGCAGGTCTAGGCGTCGAAGCGAAGTACCCTTGTAGCCGTTAGCAGAGTGGCGTTAGGCGTTGATTGGGGAAAACTGACGTAAGAAAGCGACAGCAACTAGGCGAGGAAACTGACGGACAAATCACCGGCCGGGAAGCTAGTAAGCGGCTAGATCTGTGTGGTTTTGATGCAACCACACGAAATCTCCGAATTCGAAGCAGAGAGCAGCTCTTGGGAAAGATCTGAACGGGCGTATCGACAAGACTTCTCATTCAGGAATCTCAGAACCTATCCACGTTGGGAATCAAACCAGAGGACACCCTCTCTTGAATTTCCGTGCTACCATTACAACACAACAACTGGACCTCCAGTCCACCGTACTCTCCTCAGACAAGGTGACGGCAAGGATTTACCATACTTAGTAAACACCTTGTTCGATCTCAATATCACCGAGATCCACAATCAGGCGATCCTTGACGATAAGATCTCAAGGCTCACCCAGTACCTGACAACAAAAGTCGGTTCCCTACCGACGATCCCGGAGGATTCGCCCCTCCTGGACCAAATAGCCCTTTCCCTAGATCTACAGGCCCTCAAGGCAGATCTGAAGGAAATCAAGGCCACACAGTCAGCACAAAAGCTAGCCTTCTCACAACTGCAGGAGGCAGTCCAGCTTTTTATCGCTCGGGAGAACGATCCCAAGCCGATCGAGGCAGCTACTGCACAAGTAGCCGAACAGCTGAGGAAGCAACTCATCGAGGTCAAGACAGTCCTCGAAGAGACCAAGAAGATCGCCAGGTCCTTATCCCCCGACGGATGAACCCTAGGTGGCAGGAAACTGCAGCCAAGGAAACCTATATCAAAGCGATCCAAGCAACCGCTTCCCTCACCTCCAACGGCACCGGCCAAGGCTTCATTGAGCCCCACACCTACACCGGAGGACAGTTATCTACCAATCTAGCCAAGCAAAACAACACCATAATCGAATTATTGGTGCAAGTGCTAGAAAAGAATCTCGACCTTGAACGGGCCGTAGCCAACCTCACAGGTCAA GTCACTAGGCTCGAAAAAGCCGTAGCAGACAAAGAAGCAGTCAAGCTTCCGGAAAAAGTCCTAGAAGACCTAACCAAGGAATTTGGAAAGGTTAACTTAGGCAAAGGAAAGGGGAAAAAAAGGAAAGTCTCAAGCCGAGACAGAACTTCTACGTGTGGAAAACCCCTACACTCAATACAATGAGCAGAAGCCACACAAGGCTCCAAGCGCCCCCCGCAACTGAAAGGGCCACAAGCAGCTCGGATTCAGGGACACCAACCCTGGAAGACCAGATCCGAGGATACAGGAGATCTGCACGCATGAGGCATCAAGCGCAGCAGAGACTACGGAGGACCTTCGGAAGGGACTTCAGAAACACGATCGAGAGACAACTCGATCCTGATGCAGAGCTCTCCCTCAGTCGAAGGAGAAGAGCCAACTTGGTACCGGCGGAAGTGCTCTACGCACACAACGGACAAGAACCGGTCAACCGAGTATACGAGCACTACAGTGAGCTCAGCGCTCATGTGGTAGACAGGCAGCAGGACTTCCGATTTATAGAGGAAGCGTCCTATCAAAGGCTGACCAGAGAAGGAATGCAGTTCATCCACGTAGGTATGGCGATGGTCAGGATACAAATGCTGCACAGGACAGATGCGGGTATATCCGCACTAGTGGTGTTCCGAGACACCCGATGGAGCGATGACAGGCAGGTCATCGGTAGTATGTCAGTCGATATGACACGAGGAGCGCAGTTGGTATACATCATACCCAACGCCATGATGTCGATTCATGACTTCTACAACCGGATTCAGGTCAGCATACAGACCAGAGGATACGGCACAGGCTGGGAAGGAGGCGACAGCAACATGATTATCACAAGATCATTAGTCGGCCGACTCACAAACACCAGCATCACAAGCTTCGAGTACAGGATAGACAACGTGACAGACTACCTAGCCAGCAACGGCGTAGCCTGCATTCCCGGACAGAAGTGGTCCGTGGCAAACAGGTCTGGAGAATGGGAACTCCAGCCAAGCAGGATAGCAGCACCACTTGCAGTCCCCACAGATGCCAGGCTAAGACAGAACCCAAACGGCAACATCAGCCTGAGGTTCACGGATTTCCGCGACCAAAGGATCGTGGAAGAAGGAGAGACATCCGAGCCAGAAGGAAGACCGGAGACGAAGGAGGATGAGAGCACGCACTACGTGCTTATGTTCAAACACTCCAGCCCTAGGTGGGATACGCTCGGACAGCCCAGCGGTAAATACGACTACATGGTCCGATATGACGCACCAGAACCGACCACATGGCCAACAACAAACAGAGGATGGGACGATGACCCTCCTAAGCCACCAAGCCCAAAAGGATCCTACGAGGTAAGCCTAAGAGGCGAAAAGAAGCTGAAAGAAAAGGAACTCGCAGAGTTCACCCCAGAGACAGATCTGGTCAGCCAGTGGCTAAACCAACTCTCGAACTCGGCACACAACAGTGGAGCTTCAAGCTCAGACGATGAGCCAAAGTTCGATGAGGCAGACGACGAGGACGACGTCTACAATCAGAAAACCTGGGAGAAGGAAGACCAGGAAAAGAGGGAGCTGGAACTCCAAGGGTGGAAACCCACAGGGAGGCCAGGCCTCTATGAAATGATCCCTGAGCAAGAAGAAGAAGTCTACCTCAGGTACGAGGCAGAGGACGAAGAGGAGGATCAGGAGTTGCAGGTCATAGGAGCCGCAACAATGGATGAGCCAGAGATGGAATACCCAACCAGGCTCGAAAAAGTAATGGGCAAACTCAAAAACGTAAGCATGGAGAAGCTGTTCCCAGTGAGCGGGATGGATAGCGAATCCAGCATAACAGGAGGTGGTGGAGGTTTTATACCACCAAGCCCAGTACCGGGAGCACAAGGATACCCCCCAGCAACAACATCCACTATGTCCACCATTGGACCAGCAGACATGCAGGGATGGGGAGGGAGAGTGCCCAGAAGTAGGTCACCATTAGGGTATGGCAGACCACAACAGCCATGGTCACTACCATCAGCACAGTCAGACAACGGCTGCATGCTAGTCCTTCCACAGGACTTCACCCTAATACCGGATGTCATCAACCGATGGGAATCCATAACAGTCAACCTCATCAACAAAATGATGTTTGACTCCCTGCAGGACAAGGCCGACTACGTCGAGAACCTTCTTGGAGAACGAGAAAAGGAGACGTGGATGACATGGCGGATGCAATATGAAGAGGAGTATAAACAACTCCTTACGATGAGCGGAGATGTGAGGAATATCACAGCCGCAGTCAAGCGGGTCTTCGGAGTACACGATCCGCATACAGGATCAGTCCACATCCAGAATCAAGCATATGCAGAGCTCGAAAGGCTCTACTGCAAACGGACGGACGACGTGATCCCCTTCCTATACGACTACTATCAGCTAGCAGCAAAATCGGGAAGGATGTGGCTCGGACCCGAGCTATCAGAGAAGCTGTTCAGAAAGCTTCCACCTGAGATAGGACCTACCATCGAGCAGGCCTATAAGGATCGATATCCAGGGCTGACCATTGGAGTTTTGGCCAGAGCAAACTTCATCCTGGAATATCTACAGAACGTGTGCAAGCAGGCAGCACTGCAGCGTTCGCTAAAAAGCCTGAGCTTCTGCAGGAATATGCCGGTCCCCGGATACTATGAGAAGAAGCAATATGGTATCAGAAAGGCTAAAACCTACAAAGGTAAGCCTCATCCGACCCACGTTAAGGTGATAAAAAACAAGTACAAGCATACGCAGGGGAAGAAATGCAAGTGCTACTTGTGTGGGATCGAAGGTCACTATGCCCGAGAGTGCCCAAAGAAGGTGGTCAAACCACAACGAGCGGCCTATTTCAATGGCATGGGACTAGACGATAACTGGGATGTAGTCTCGGTCGAACCAGGAGAAGAAGACGACGACGAGATCTGCAGCATCTCAGAAGGAGAAAACACTGGCGGAATGCACGAACTTATGGCATTCAAAACACAACTCCCTTATCCAGTGGAGTATGAAGCCAGCACACCACAGTTCCTGATGCCATGGACACAGGTACCTGTGGAAAAGAGCGACAAACCTTCCTGGAGAAGACGGAAGGATATCTCACAAGTCCAGAAGGACTGCACACACACCTGGAGTGACACCCAGGAAGTGCCTATCAGCGACAGGGTTTGCAGCATCTGTAGTGACGAAACACCTCACGGTAGAAGAGTCACCTGCACTACATGCAACATCAACCTCTGTCCGATATGTGCAAGGATGGACTATGGGATCATGCTGATAGCAGCAAAAGACACCAAGAGCGCAGCACATTGGCAGTACCAAAACAAGGATGAGCTCATACAGCACCTGTACGAACACAACGCTTTTCTCACCAGGAAAGTAGCAGAGCTCACTAGCCAGCTGCAGGAATTCCACAACCGCAGGCCTGAAGACCTGATCAGCCTAGCGGATGACCTGGAGGACGTGTCCATCCTGGACAACGCCTCAAACAGGGGGAAGGAGGAGAAGGAATTGTTCCAATTTGGAACTACAATTCCCATCGACCACATACAAAACCTTGAAAATGTGGCCAAGATCATAGAAAAGTGGAAAGATACCCCCAGGGTCGTAATCAAGGAAACACCAGAAAGCAGTACCAGTAACACCATCGGAGCACTTCTAGCTGAGGAAGGAATCGAAGAACTGGCTGCAGCGGTTGACACCGCCTATACCGAGATGCCAAAAGGAGGCCTCAACAAACTTTACAACACAATTGTGGAGTTCGTCATACCTCAGGAAAAGGGAGCACCCACGAAGTTCAGAGTTCGTGCTGTGATAGATACTGGATGCACCTGCACGTGCATTAACAGTAAAAAGGTTCCCAAGGAGGCACTCGAAGAAGCGAAGTACCAGATGAACTTCGCAGGGGTTAACTCAACAGGGGAGACCAAGCTGAAAATGAAGAACGGGAAGATGATCGTGTCAGGCAGCGACTTCTACACCCCGTATATAGCAGCTTTCCCAATGGAGTTGCCAGACGTAGACATGCTGATTGGCTGCAACTTCTTGCGAGCCATGAAGGGAGGCGTAAGACTCGAAGGAACGGAAGTGACTATCTACAAGAAAGTCACCACAATCCAAACAACCCTAGAGCCACAGAAGATATCCCTCCTCCGAGCAGAGGCTGAAGTCGGAGAAGAACTAGAGCGCATGTACTACGCCAATGACTATTCCGAGGAAGGAATAAGTCGGCTGAAGAACCACAGGCTGCTGCAGGAACTCAGAGAACAAGGGTACATTGGTGAAGAGCCAATGAGACACTGGGCAAAGAACGGCATCAAATGCAAGCTGGATATC AAGAATCCAGACATAGTCATCAGCAGCAAACCGCCTGACTCTGTATCAAAAGAGACGAAAGCCCAATACCAAAGGCATATAGATGCCCTGCTCAAAATCGGAGTAATCCAGCCCAGCAAGAGCAAACACAGGACGGCGGCTTTTATCACACACTCGGGTACGTCAATTGACCCGATTACCAAGAAAGAAGTCAGAGGGAAAGAACGGATGGTATTCGACTACCGAAGTCTCAACGACAATACCCACAAAGATCAGTACACACTGCCGGGTATCAATACCATCATATCCGCGATTGGCAACGCTAAAATATTTAGCAAGTTCGATCTAAAGTCCGGATTCCATCAGGTGCTCATGGACGAAGAATCCATACCATGGACGGCTTTCGTAACGCCAGTCGGATTCTATGAATGGAAGGTCATGCCCTTTGGCCTTGCCAATGCTCCAGCTGTCTTCCAAAGGAAGATGGACCAATGCTTCGCTGGAACTTCGGAATTCATCGCAGTCTACATCGATGACATCCTGGTGTTCAGCAAAACCCTAAAGGAGCACGAGAAACACCTTAGCATCATGCTAGGGATATGCCGTGATAACGGTTTGGTTTTATCGCCCAGCAAAATGAAGTTGGCCGCCACAGAGATAGACTTCCTTGGCGCCACCATAGGCGATGGAAGGATCAAGCTCCAGCCTCACATCATAAAGAAGATAGCCGAGGTGGATGACGAATCCCTGAAAACCCTCAAAGGGTTACGAAGCTGGTTGGGAGTGCTCAACTACGCGCGCAACTACATCCCAAAGTGTGGCACACTGTTAGGCCCACTATACAGCAAGACCAGCGAGCATGGTGATCGTAGGTGGCACGCGTCTGATTGGGCCTTAGTCAAAAGAATTAAGGGCCTGGTCCAAAACCTCCCAGACCTAAAACTCCCCACGGAAGAGGCATACATGATCATTGAGACTGATGGATGCATGGAGGGCTGGGGAGGAGTCTGCAAATGGAAGCCCATGAAGGCAGACTCAGCAAGCAAGGAAGAAATCTGCGCTTACGCCAGTGGTAAATTCCCCACGGTAAAATCAACAATAGACGCAGAAATCTTCGCAGTTATGGAGTCCTTGGAAAAATTCAAGATTTTTTACATGAACAAGGACGAGGTCACTATCAGGACTGATTGTCAAGCAATAATCACCTTCTACGAGAAGCTGAATGCAAAGAAACCTTCGAGGGTAAGGTGGCTTGCCTTTTGCGATTATATAACGAACTCCGGGGTAAGAATGAAGTTCGAACATATAAAAGGTAAAGACAACCAGCTCGCAGATAACCTCAGCCGTCTCACACAACTGATTACATTTGTGAAATGGCTTCCAACCGAGCTCAAGGACCTCGCGGCAGAACTAACCAGGAAAGACGACGGGACGCCCGCGAAGAAGGAAGTGCAGGAGGAAATCTCCTGTTTTCTCGAAGCTGCCCTCCGCCGAGCCAAGAGATCCGTGACTACTCACCAATCCGAGCCACGCCATGTACTATGGCAGAAATGGCAGAATCCAGAAGGCTGGCTCTACTGCGACGAGAGGAGATCTTCAACAGCCTTGCCCAACACATCAGCGACACGGTCTTCATCACCGGAGTCGACCTTGCGGCAGCAAAGGCCAGAGCAACCAGGGACAACTGGTATGCTGACATCACACCAACACTGGAACGACGAGCCACCGCAGCATGGAAGCTCATGGCCGCTTACGAGGAATTCGCCACGTGTAAGGATGTGAACGTTTAGTGAAGCGACGTCAGCAATGACTTCACAATCGCCCAAGTGCGTCACTGCTTACGCTTGGGAACTTATCTTTTAGTGTCGGTAGCATCTTCTAGCTGCCATACTTTATTGTAAGTGCGCCGATAGTGCGCTGAGTCATAGTGATAAGGAATCTTATCACCTTATCGTCCTTTCTTAGCTTTAGTAGCTGTAAAGACGAACTTATTAGCTGTCGATGGGGCCCAGAAAGCGCACCCGAGCTGATATTTTCTCTCTTTCTGCTAAGCCCTCCCCCTATATAAGGGAGAAGAGTTTGAAGGCTTAGGCACAGAGCAATCTCTCTAGCCAACCTTTCTCTTGAGTTGTATTAAAACATTCAAGTGAAATAAAGACTTGTTCATCTTTTTCCGCATATCTCTGAGTTTTTATGAGTTCTTAAGTGTTCGAAAGCGCACTTTTGAAATTAGATCCATGTTTTTCGGACCCCATTC

**Supplemental table.**
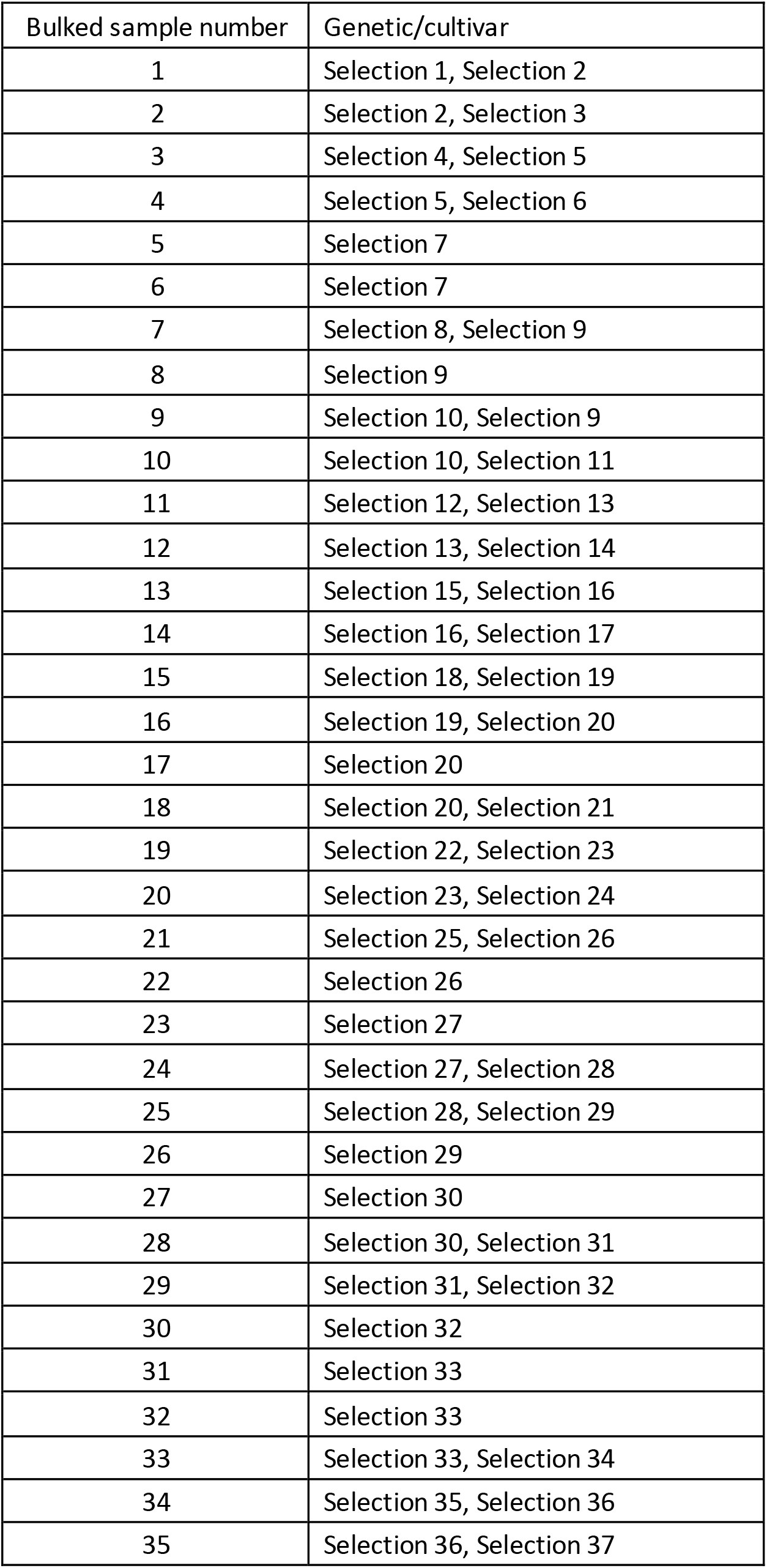
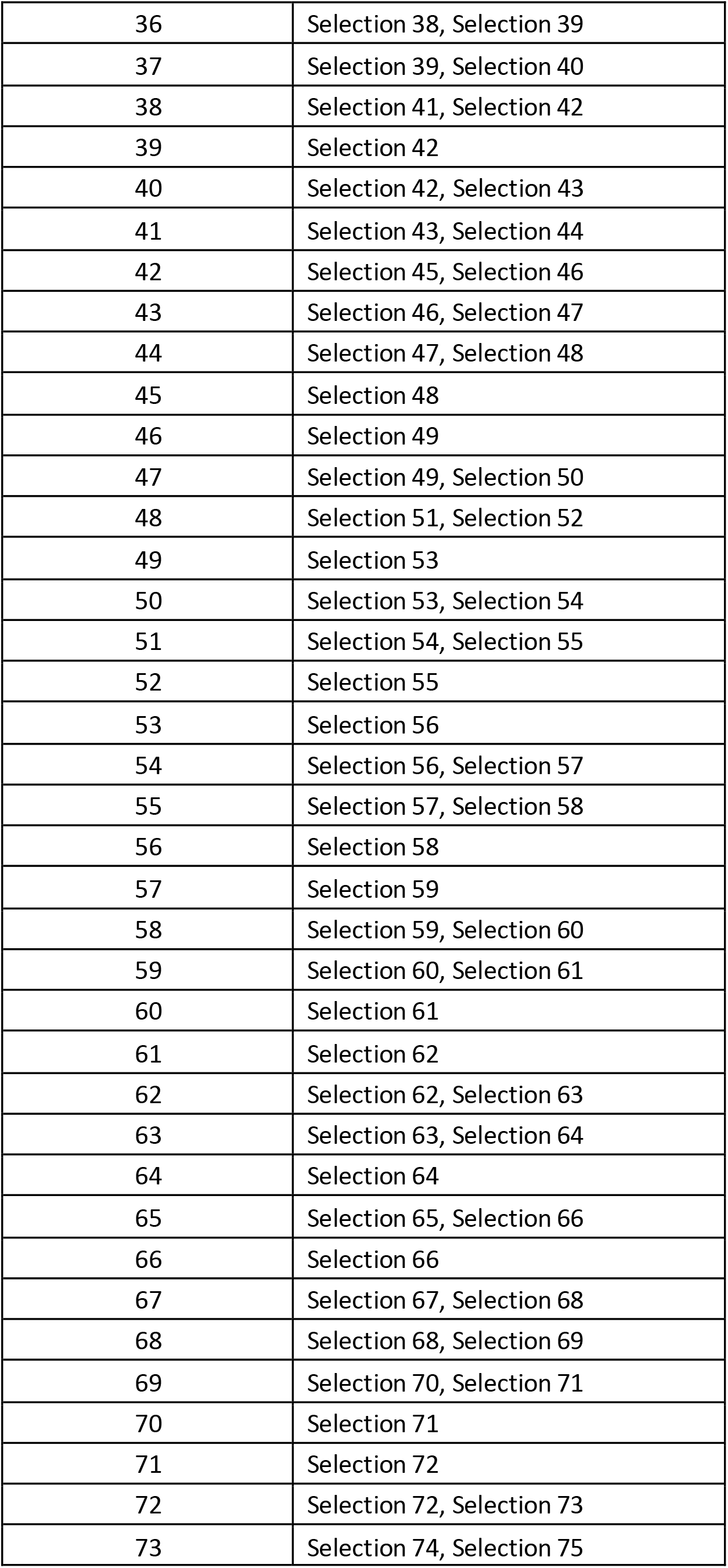
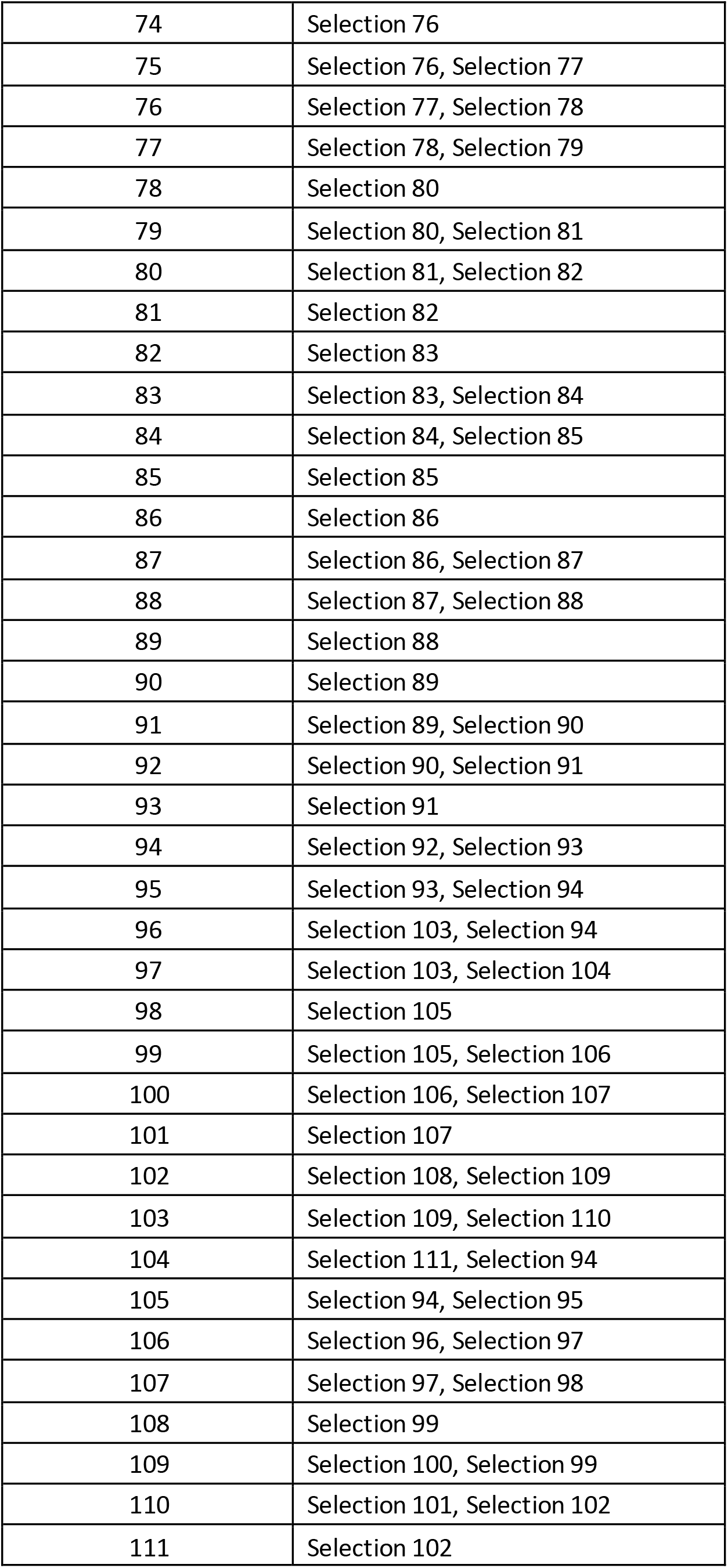
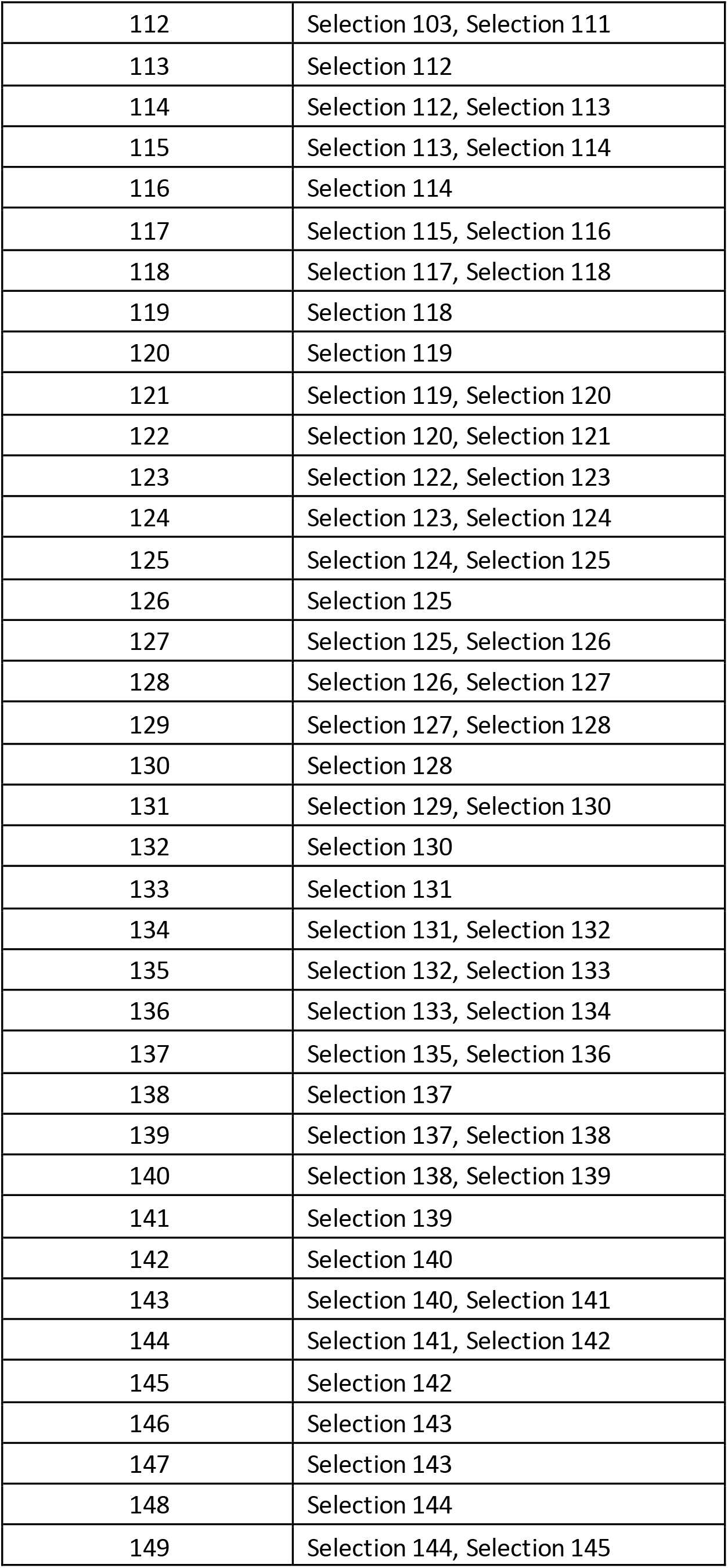
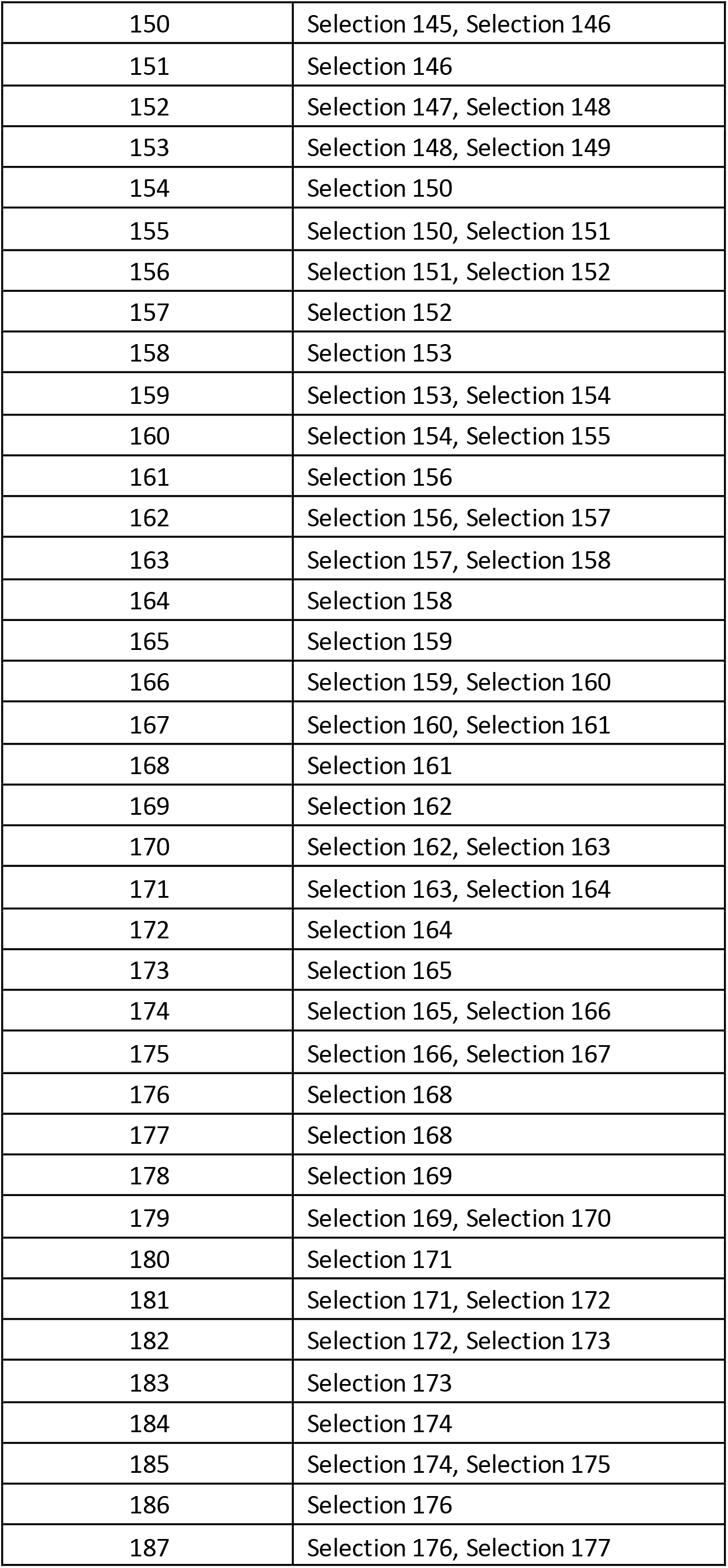
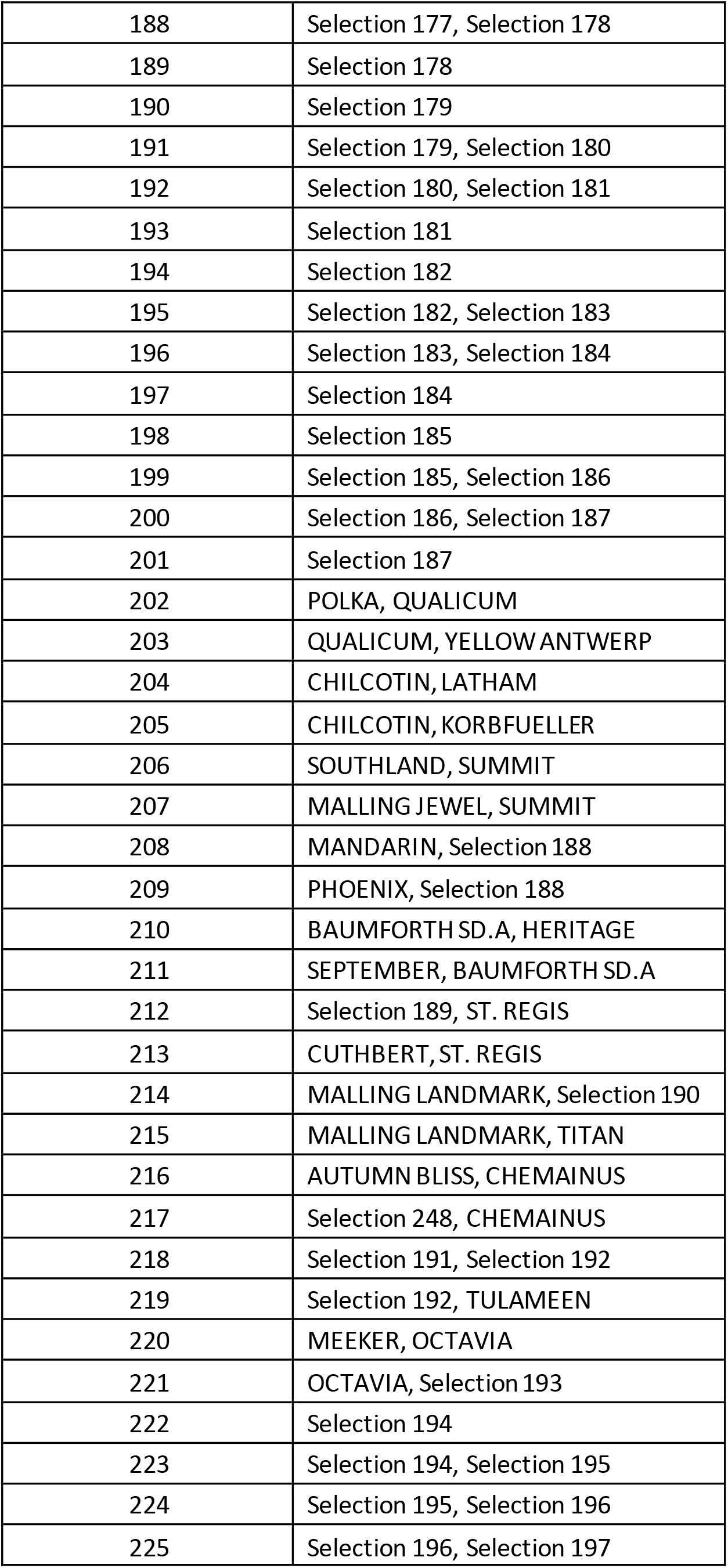
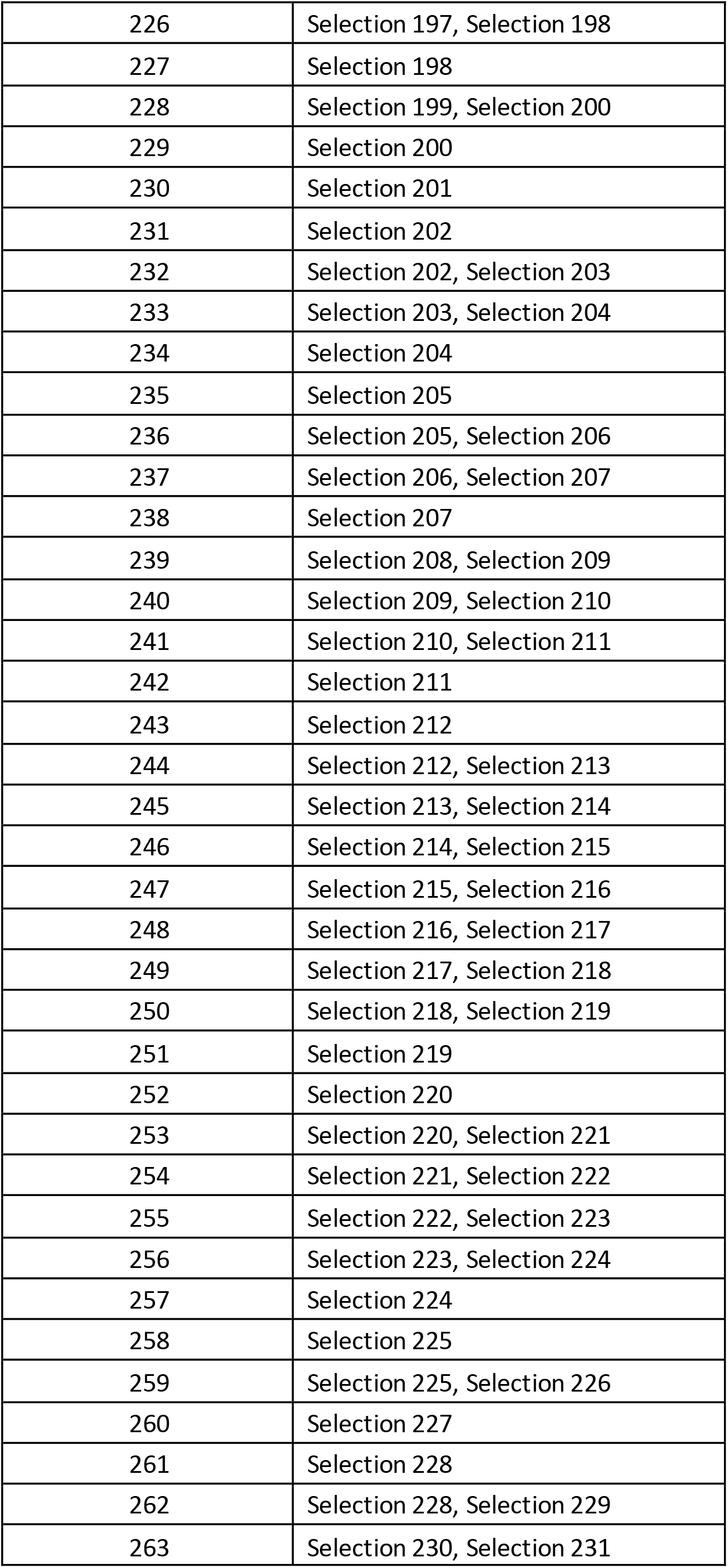
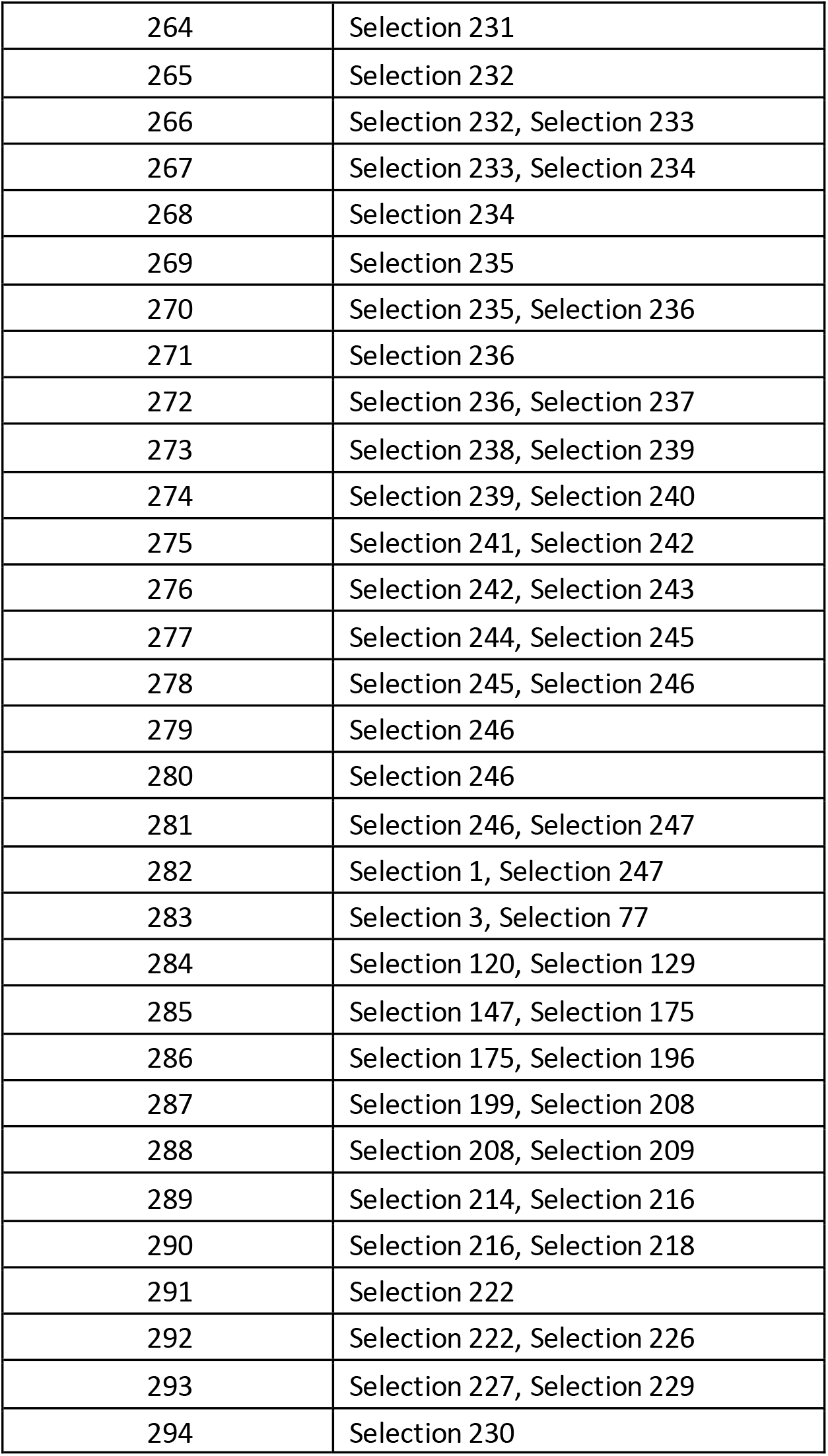
271 public and proprietary genetics were bulked into 294 samples and used in validation of epiRYNV-BS detectionassay.

**Supplemental figure.**
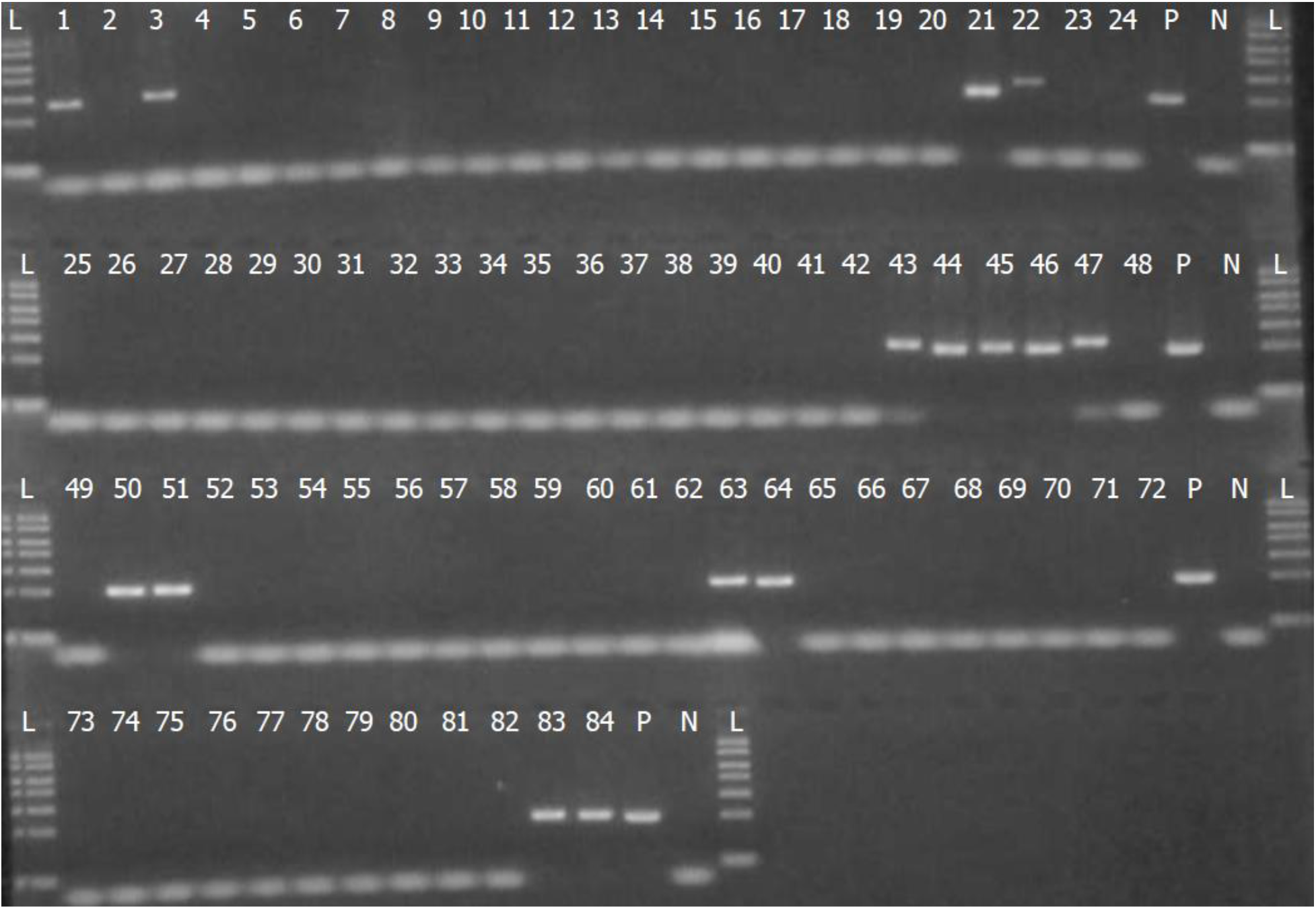
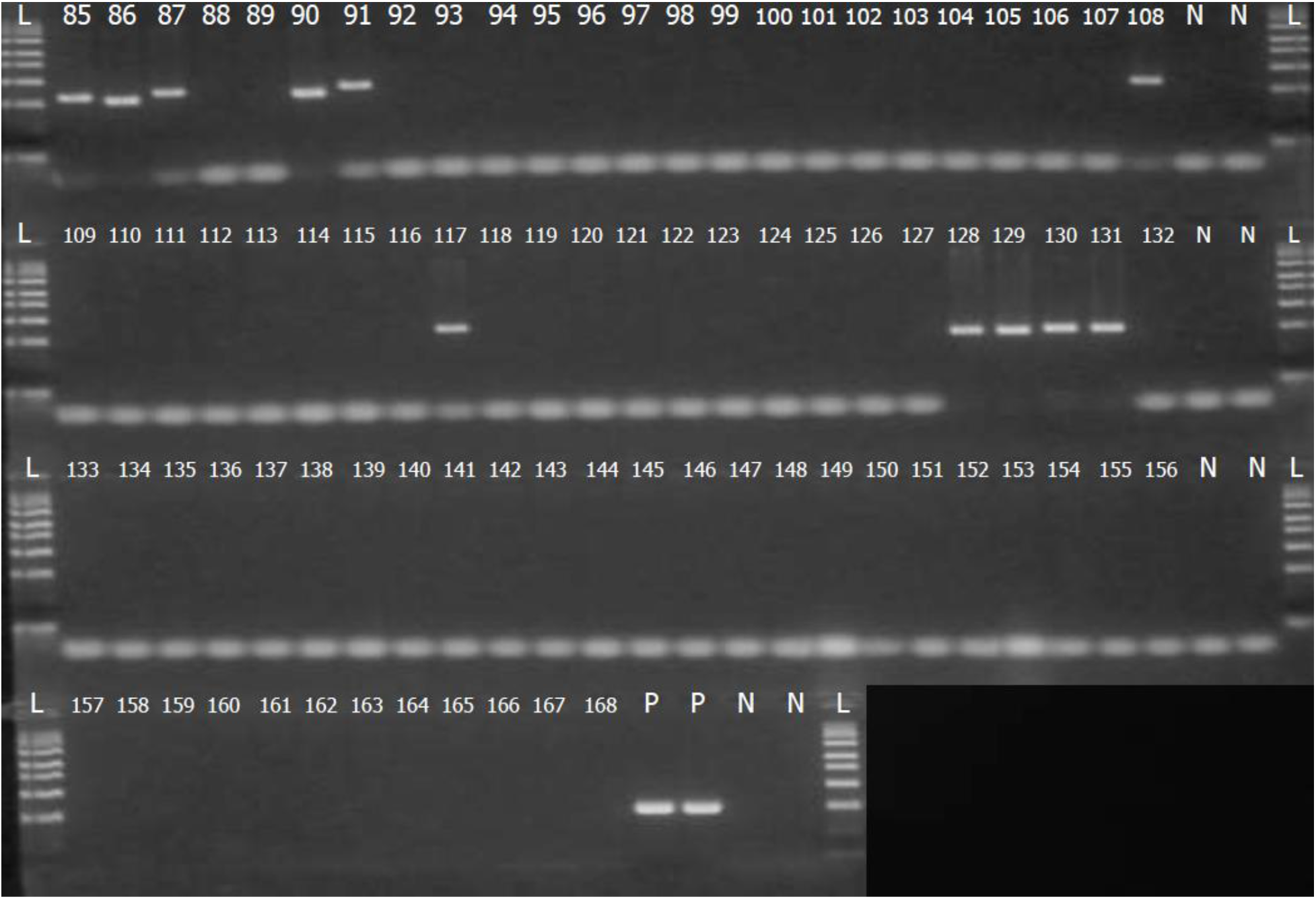
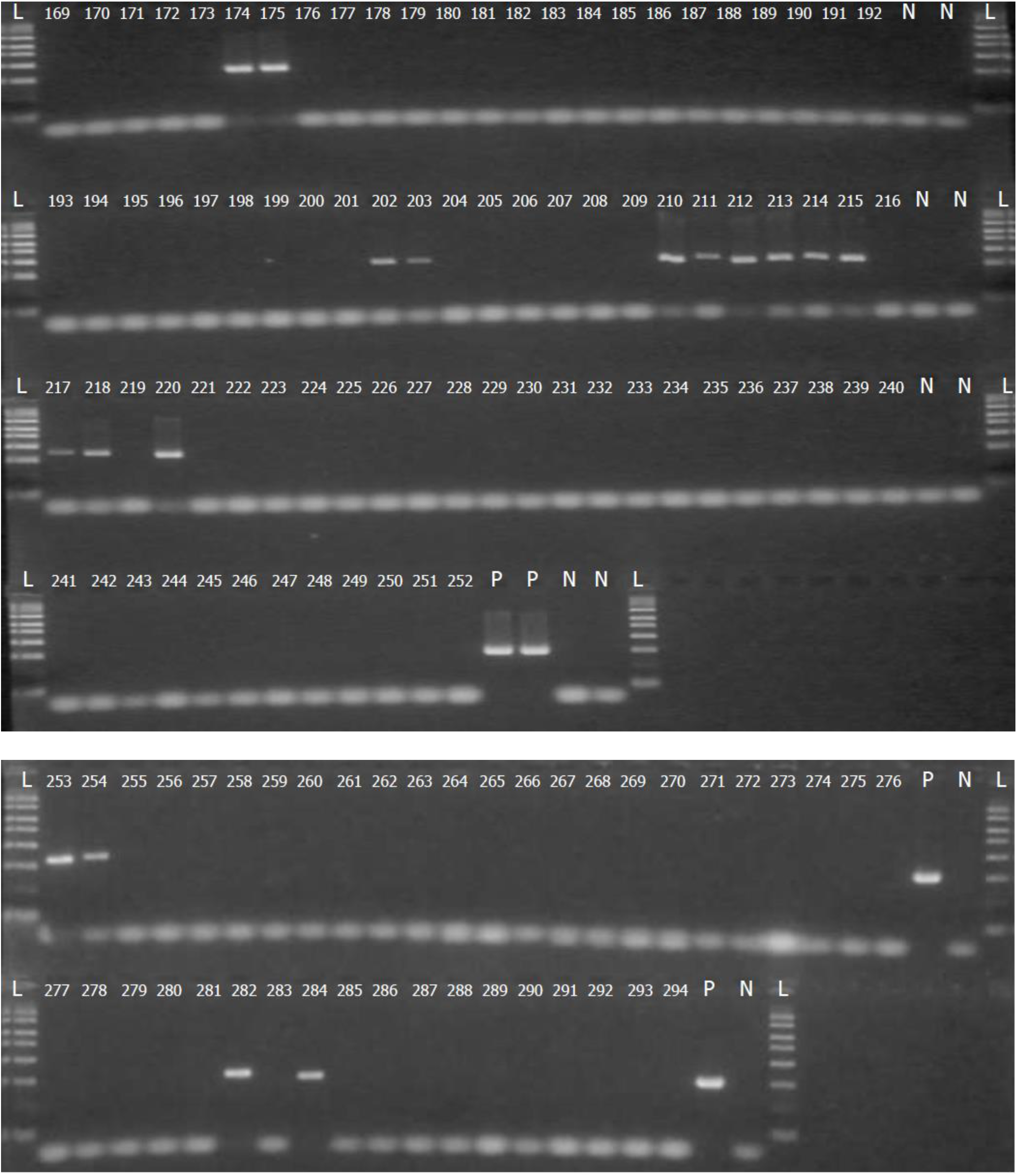
epiRYNV-BS detection using RYNV6-F/R PCR assay. 271 public and proprietary genetics were bulked into 294 samples. Off-target was observed in 43 cases.

